# Th17-skewed inflammation due to genetic deficiency of a cadherin stress sensor

**DOI:** 10.1101/2020.12.01.406587

**Authors:** Lisa M Godsel, Quinn R Roth-Carter, Jennifer L Koetsier, Lam C Tsoi, Joshua A Broussard, Gillian N Fitz, Sarah M Lloyd, Junghun Kweon, Amber L Huffine, Hope E Burks, Marihan Hegazy, Saki Amagai, Paul W. Harms, Jodi L Johnson, Gloria Urciuoli, Lynn T. Doglio, William R Swindell, Rajeshwar Awatramani, Eli Sprecher, Xiaomin Bao, Eran Cohen-Barak, Caterina Missero, Johann E Gudjonsson, Kathleen J Green

## Abstract

Desmoglein 1 (Dsg1) is a cadherin restricted to stratified tissues of terrestrial vertebrates, which serve as essential physical and immune barriers. Dsg1’s importance in epidermal integrity is underscored by genetic, autoimmune and bacterial toxin-mediated disorders interfering with Dsg1 function. Dsg1 loss-of-function mutations in humans result not only in skin lesions, but also multiple allergies, and isolated patient keratinocytes exhibit increased pro-allergic cytokine expression. However, the mechanism by which genetic deficiency of Dsg1 causes chronic inflammation is unknown. To determine the systemic response to Dsg1 loss, we deleted the three tandem Dsg1 genes in mice using CRISPR/Cas9. Whole transcriptome analysis of E18.5 *Dsg1*^−/−^ skin showed changes consistent with the observed aberrant differentiation and barrier impairment. Comparing epidermal transcriptomes from E18.5 Dsg1-deficient mice and humans with Dsg1 mutations revealed a shared psoriatic-like IL-17-skewed inflammatory signature and less so a pro-allergic IL-4/13 signature. Although the impaired intercellular adhesion observed in *Dsg1*^−/−^ mice resembles that resulting from autoimmune anti-Dsg1 pemphigus foliaceus antibodies, transcriptomic analysis of pemphigus skin lesions lacks a prominent IL-17 signature. Thus, beyond impairing the physical barrier, chronic loss of Dsg1 function through gene mutation results in a psoriatic-like inflammatory signature before birth, possibly predisposing to skin inflammation.

## Introduction

The multi-layered epidermal barrier is made up of interdependent microbiome, chemical, physical and immune components. These components work together to protect against water loss, physical insults, and infection (1, 2). The asymmetric distribution of membrane proteins along the apical to basal axis of simple epithelia ensures that epithelial barrier and transport functions are properly regulated. However, in multi-layered epithelia such as the epidermis, architectural features are polarized along the entire apical to basal axis of the stratified epithelium. Disorganization or loss of these polarized features disrupts barrier function of the epidermis and causes a range of cutaneous diseases (3).

Among the most polarized molecules in the epidermis are desmosomal cadherins, desmogleins (Dsgs) and desmocollins (Dscs) (4, 5). These transmembrane cell-cell adhesion molecules cooperate with plakins and armadillo proteins to anchor intermediate filaments (IF) to the desmosome, providing tissues with tensile strength. Mutations, bacterial toxins, pemphigus autoimmune antibodies and dysregulated expression of these cadherins cause a range of mild to potentially lethal disorders including keratodermas, blistering diseases and cancer in humans and mouse models (5–9).

Among the recently identified disorders caused by mutations in desmosome molecules is severe dermatitis, multiple allergies and metabolic wasting (SAM) syndrome (10–15). This disorder was initially identified in patients harboring bi-allelic mutations in desmoglein 1 (Dsg1), resulting in a reduction of Dsg1 expression and/or failure to accumulate at the plasma membrane. SAM syndrome patients exhibit abnormal epidermal differentiation, recurrent skin infections and severe allergies. While the loss of barrier forming roles of Dsg1 may contribute to these symptoms, isolated patient keratinocytes exhibit cell autonomous production of cytokines, including the pro-allergic cytokines thymic stromal lymphopoietin (*TSLP*), *IL-5* and tumor necrosis factor (*TNF*) (10, 14). Further, knockdown of Dsg1 in normal human keratinocytes *in vitro* was sufficient to induce the expression of pro-inflammatory cytokines *IL-1β*, *IL-6*, *IL-8*, *CXCL1* and *TNF* (14, 16). Combined with the observation that Dsg1 is downregulated in response to environmental stress, such as exposure to UV, these observations raise the possibility that Dsg1 is a stress sensor that governs the expression of cytokines independently of its role in maintaining tissue integrity and barrier function (10, 16–18).

Functions for desmosomal cadherins that transcend adhesive roles have also recently emerged. For instance, Dsg1 promotes keratinocyte differentiation as cells transit from the basal layer through attenuation of Ras-Raf signaling to Erk, which requires Dsg1 binding to ErbB2 Interacting Protein, Erbin (19, 20). Attenuation of this pathway by Dsg1 facilitates differentiation without inhibiting basal cell proliferation. In addition to its role in harnessing Erk signaling, we showed that Dsg1 remodels the cortical actin cytoskeleton to temporarily reduce tension in basal cells, which is necessary for promoting stratification in human epidermal organotypic cultures (21). Thus, Dsg1 coordinates a transcriptional and morphological program of epidermal differentiation and morphogenesis in an *in vitro* human model of epidermal morphogenesis. Progress in elucidating adhesion-dependent and -independent functions of Dsg1 *in vivo* has been hampered by the lack of a fully characterized animal model.

Here we report results from an animal model in which the tandemly arrayed *Dsg1 a, b* and *c* genes were removed using CRISPR/Cas9 mediated gene editing. In addition to analysis of severe adhesion and barrier defects in Dsg1 deficient animals, comparisons between whole transcriptome profiles of the embryonic skin of Dsg1-deficient animals and skin from SAM syndrome patients were performed. Analyses of these profiles revealed shared changes in epidermal keratinization, keratinocyte differentiation and skin development pathways and an increase in inflammatory response pathways. We compared these data sets with patient cohorts from two common inflammatory disorders, atopic dermatitis, largely a Th2-dependent disorder, and psoriasis, largely a Th17-dependent disorder (22). The previously reported increase in SAM syndrome patients’ keratinocyte cytokines was weighted toward a Th2 response (10, 14); however, the transcriptome analyses of both Dsg1 deficient animals and SAM syndrome biopsies revealed an inflammatory response that was skewed towards Th17 and up-regulation of IL-36 response genes. Both Th17 and IL-36 play crucial roles in psoriatic inflammation (22, 23). Consistent with this observation, gene signatures from both the Dsg1-deficient animals and lesional skin from SAM syndrome patients showed significant similarity to gene signatures from a cohort of psoriasis patients and less so to a cohort of atopic dermatitis patients. Interestingly, while morphological features of Dsg1-deficient epidermis resemble those found in human pemphigus foliaceus (PF) patients, transcriptomic analysis of PF lesions lacked the Th17 skewing exhibited by targeting the Dsg1 gene. As the IL-17/23 skewed gene signature is present before birth in Dsg1 null embryos, genetically induced loss of Dsg1 could predispose individuals to skin inflammation.

## Results

### *Dsg1*^−/−^ mice exhibit loss of cell-cell adhesion and severe epidermal peeling in spite of increased Dsc1 and Dsg3

Dsg1 stands out among the other desmosomal cadherins as having three genes instead of one in the mouse, *Dsg1a*, *b* and *c*, located on chromosome 18 (24, 25). The functional differences among these three genes in various tissues are not well understood (25). qRT-PCR of *Dsg1^+/+^* mouse tissue revealed that only *Dsg1a* and *b* genes are expressed in skin, as well as the esophagus, tongue and forestomach, while *Dsg1c* expression was found only in the liver (Supplemental Figure 1A). To generate a Dsg1 knockout mouse, two different methods were employed utilizing CRISPR/Cas9 technology (Supplemental Figure 1B). In one model exon 2 was deleted in the *Dsg1a* and *b* genes to mimic a previously characterized SAM syndrome mutation using gRNAs flanking exon 2 in each gene (10). Loss of exon 2, which was confirmed by PCR on genomic DNA, resulted in perinatal lethality of the chimeric animals. While we were unable to generate mice with germline transmission, blisters were observed in the skin of chimeric animals and histological analyses revealed areas of Dsg1 loss associated with epidermal blistering at the junction between the granular and cornified layers (Supplemental Figure 1C, D). We also observed a reduction in total Dsg1 in the skin of chimeric animals by immunoblot, indicating that deletion of exon 2 causes reduced stability of Dsg1 (Supplemental Figure 1E).

In the second model, the tandemly organized *Dsg1a*, *b* and *c* gene cluster was removed by gene editing (Supplemental Figure 1B). Sequence analysis of progeny confirmed loss of the *Dsg1* gene cluster in knockout animals. A complete loss or reduction in Dsg1 protein (Figure 1A-C) and mRNA (Figure 1D, Supplemental Figure 2A) was also observed in *Dsg1*^−/−^ and *Dsg1^+/−^* animals, respectively. *Dsg1*^−/−^ animals are similar in size to their *Dsg1^+/+^* and *Dsg1^+/−^* littermates and were born at normal Mendelian ratios (103 *Dsg1*^−/−^ animals/410 total animals analyzed). *Dsg1*^−/−^ animals die within hours of birth and exhibit severe skin fragility and perinatal lethality with peeling skin (Figure 1G). Histologic analysis revealed a split between the granular and cornified layers of epidermis from *Dsg1*^−/−^, but not *Dsg1^+/+^*or *Dsg1^+/−^*, animals (n=11-17) with the stratum corneum (SC) separating completely from the tissue (Figure 1H). Increased intercellular spaces were observed in the granular layer by electron microscopy, suggesting that adhesion defects are also present in this layer (Supplemental Figure 2B).

**Figure 1.**
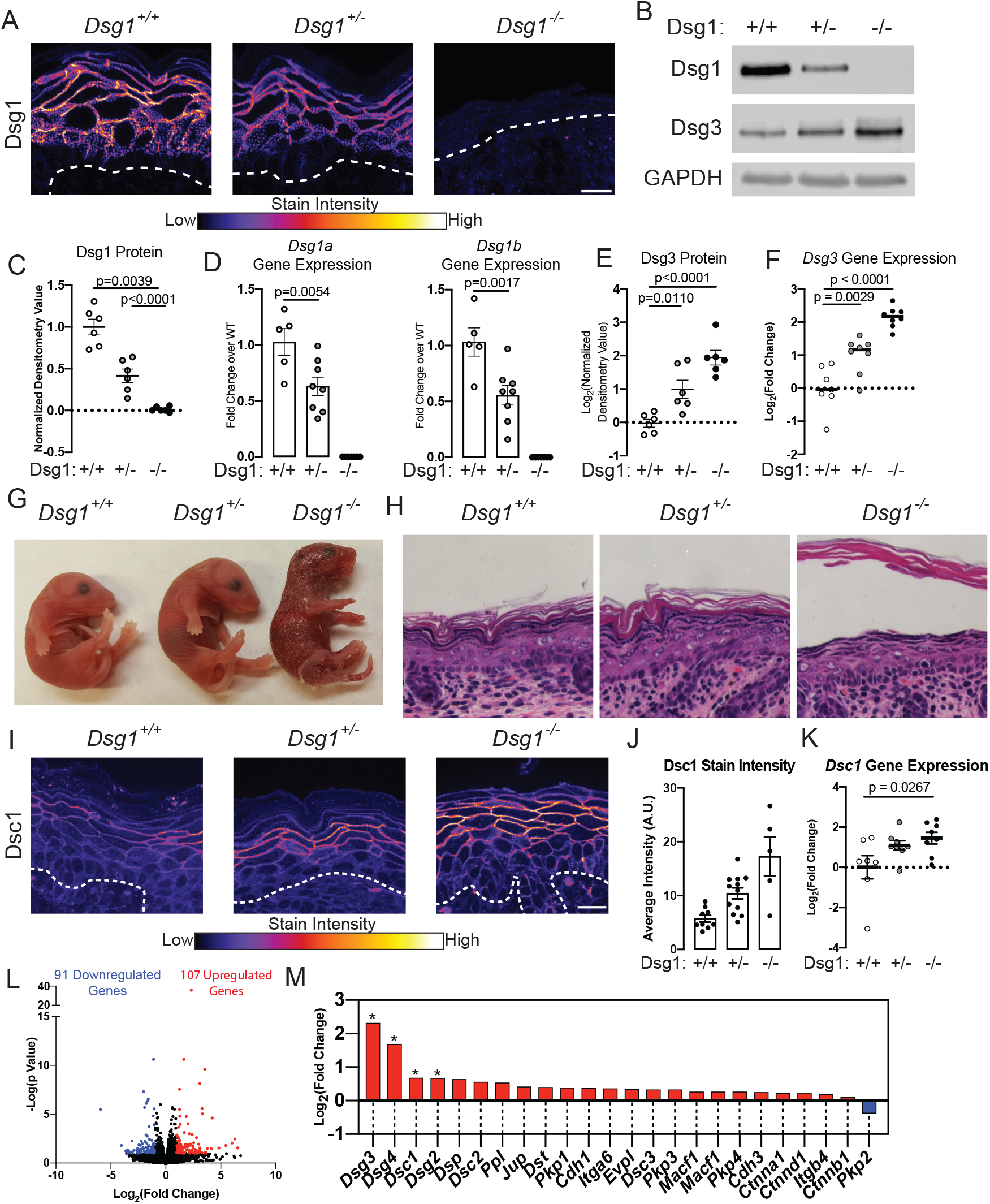
*Dsg1*^−/−^ mice exhibit defects in epidermal adhesion. A) Immunostaining for Dsg1 in E18.5 mouse skin. Dashed line indicates location of the basement membrane. Scale bar = 20 μm. B) Immunoblot for Dsg1 and Dsg3 in protein extracts from E18.5 mouse skin. GAPDH was used as a loading control. C) Quantification of Dsg1 protein from immunoblots (n = 6 mice/genotype). Densitometry values were normalized to the *Dsg1^+/+^* samples and GAPDH was used as a loading control. *Dsg1a* and *Dsg1b* gene expression in whole skin from E18.5 mice (n = 5-8/genotype). E) Quantification of Dsg3 protein from immunoblot of protein extracts from E18.5 mouse skin. Densitometry values were normalized to the *Dsg1^+/+^* samples and GAPDH was used as a loading control (n = 6/genotype). F) *Dsg3* gene expression in skin from E18.5 mice (n = 5-8/genotype). G) Images of P1 neonates. H) H & E staining of skin from E18.5 mice. I) Immunostaining for Dsc1 in E18.5 mouse skin. Dashed line represents basement membrane. Scale Bar = 20 μm. J) Staining intensity of Dsc1 in the epidermis of E18.5 mouse skin (n = 5-12). K) *Dsc1* gene expression in skin from E18.5 mice (n=6-8). L) Volcano plot of upregulated and downregulated genes from RNA-Seq analysis performed on E18.5 mouse skin. Genes are considered significantly changed if they are p<0.1 and greater than two-fold increased or decreased in *Dsg1*^−/−^ mice compared to *Dsg1^+/+^* (n = 5). M) mRNA expression levels for proteins that make up desmosomes, adherens junctions, and hemidesmosomes from the RNA-Seq data set.

Based upon the observed adhesion defects in the skin of *Dsg1*^−/−^ animals, we determined levels of mRNA and protein for other desmosomal and classic cadherins and their associated proteins in the epidermis (Supplemental Figure 2A, C, D). E-cadherin (Ecad, *Cdh1*) and P-cadherin (Pcad, *Cdh3*) gene expression was unchanged and Ecad, desmoplakin (DP, *Dsp*), and plakoglobin (PG, *Jup*) protein levels and distribution within the epidermal layers were unchanged in *Dsg1^+/−^* and *Dsg1*^−/−^ animals compared to *Dsg1^+/+^* animals (Supplemental Figure 2C, D). In contrast, the desmosomal cadherin Dsg3, which normally exhibits a reciprocal distribution pattern compared with Dsg1 (26), was increased at both total protein and gene expression levels in *Dsg1*^−/−^ animals (Figure 1B, E, F). Immunofluorescence staining showed an expanded Dsg3 distribution into the superficial spinous layers in *Dsg1*^−/−^ epidermis compared to the restricted distribution in basal layers of *Dsg1*^+/+^ epidermis (Supplemental Figure 2C). We also observed a corresponding redistribution of *Dsg3* mRNA in the *Dsg1*^−/−^ epidermis by RNAscope, with *Dsg3* gene expression restricted to the basal layers in *Dsg1^+/+^* epidermis, while *Dsg3* expression expanded into the upper layers in *Dsg1*^−/−^ epidermis (Supplemental Figure 2A). Staining intensity for Dsc1, which is normally expressed in the most suprabasal layers of the epidermis, was increased in *Dsg1*^−/−^ epidermis compared to control (Figure 1I, J), and *Dsc1* gene expression was increased in *Dsg1*^−/−^ skin (Figure 1K).

To comprehensively interrogate gene expression differences between *Dsg1^+/+^, Dsg1^+/−^* and *Dsg1*^−/−^ animals, RNA-seq whole transcriptome analysis on RNA isolated from E18.5 skin was performed. 107 significantly upregulated and 91 significantly downregulated genes (False Discovery Rate (FDR) ≤10% and |log_2_ Fold Change (FC)| ≥ 1) were observed in *Dsg1*^−/−^ animals compared to *Dsg1^+/+^* (Figure 1L). There were few changed genes in *Dsg1^+/−^* animals compared to *Dsg1^+/+^*, so all further analyses were performed using the *Dsg1*^−/−^ animals. This observation of limited changes in *Dsg1^+/−^* animals is consistent with no observable phenotype at baseline in these animals. Genes for the desmosomal cadherins *Dsg3*, *Dsg2*, and *Dsg4* were significantly increased in this RNA-seq data set (Figure 1M). *Dsc1* was modestly upregulated in the RNAseq and by qRT-PCR analysis (Figure 1K, M). We did not observe changes in expression of the desmosomal plaque components *Dsp*, *Jup* and the plakophilins or the classic cadherins, *Cdh1* and *Cdh3* (Figure 1M). While there were some differences observed between the RNA-seq and qRT-PCR data sets for significantly changed genes, all genes trended in the same direction across both datasets (Figure 1M and Supplemental Figure 2D)

### The epidermal differentiation program and barrier function are disrupted in *Dsg1*^−/−^ animals

The severe epidermal peeling and observed disruption of epidermal adhesion, along with our previous work showing that Dsg1 promotes keratinocyte differentiation *in vitro* (19), prompted us to assess differentiation and barrier functions in the *Dsg1*^−/−^ animals. Loricrin, a structural component of the cornified envelope, was decreased in *Dsg1*^−/−^ mice as observed by immunoblot and immunofluorescence, without a change in mRNA levels, suggesting that Dsg1 loss is important for the post-transcriptional regulation of loricrin expression (Figure 2A-D). Not all differentiation-associated proteins were affected, as protein levels of involucrin and transglutaminase were unchanged in the epidermis of *Dsg1*^−/−^ mice (Supplemental Figure 3A-C).

**Figure 2.**
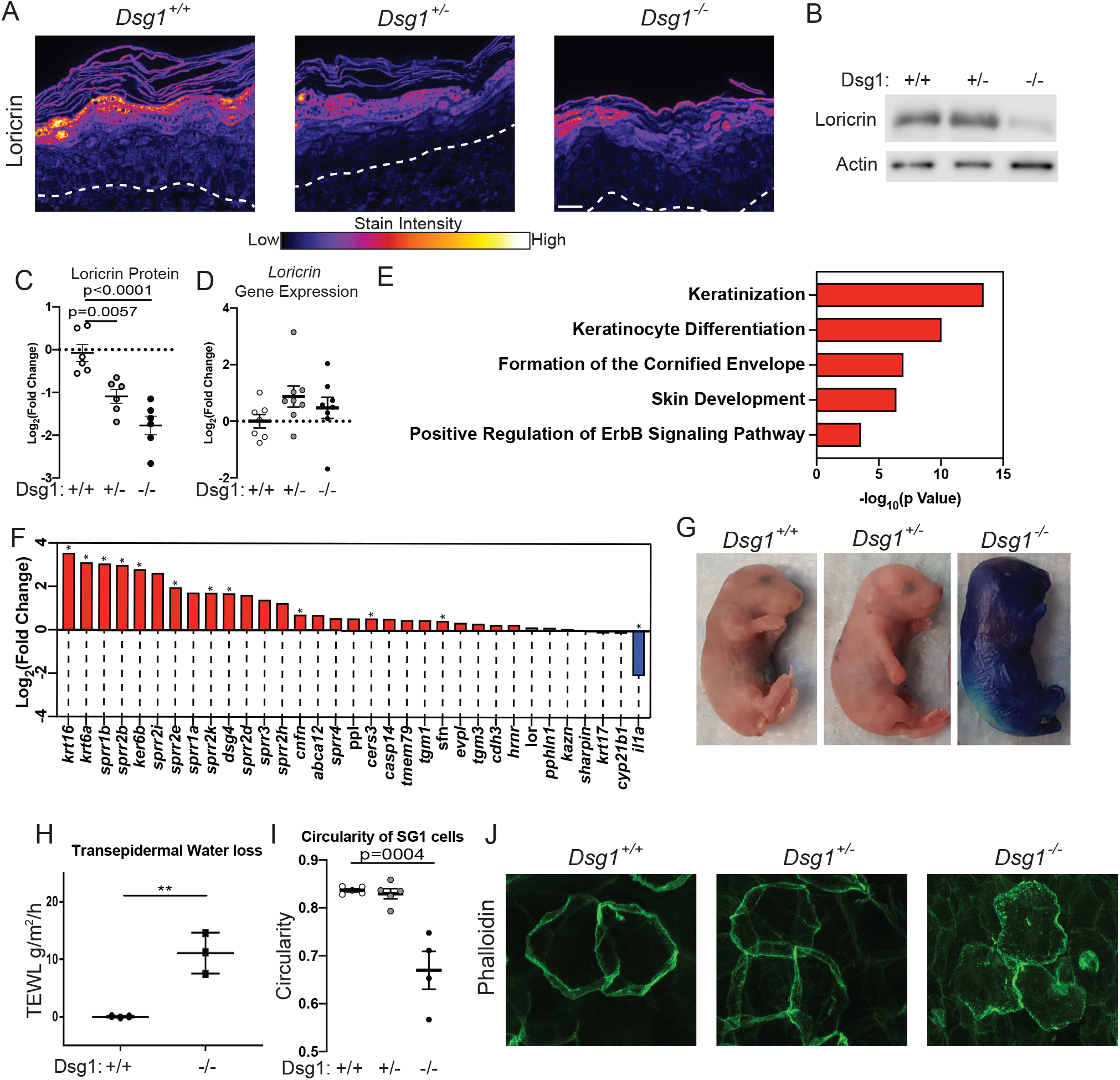
Dsg1 knockout interferes with keratinocyte differentiation and epidermal barrier function. A) Loricrin immunostaining in skin from E18.5 mice. Scale bar = 20 μm. B) Immunoblot for loricrin in protein extracts from E18.5 mouse skin. Actin was used as loading control. C) Quantification of loricrin protein from immunoblot. Densitometry values were normalized to the *Dsg1^+/+^* samples and actin was used as a loading control (n = 6/genotype). D) *Loricrin* gene expression in skin from E18.5 mice. E) Gene Ontology (GO) Biological Process terms significantly overrepresented in upregulated genes in *Dsg1*^−/−^ skin. F) Genes associated with the Keratinization GO pathway and fold change of each gene in *Dsg1*^−/−^ skin. G) Toluidine blue barrier assays performed on E18.5 embryos demonstrating an outside-in barrier defect in *Dsg1*^−/−^ animals. H) Transepidermal water loss measured in P1 pups ~5 hours after birth. I) Quantification of cell circularity in the SG1 layer in epidermal whole mounts from E18.5 mice, stained with phalloidin. J) Representative imaging of phalloidin staining of epidermal whole mounts from E18.5 mice.

To address on a more global scale whether Dsg1-deficient mice exhibit abnormalities in epidermal differentiation and barrier function, we carried out further analysis of the RNA-Seq data. Functional enrichment analysis of upregulated genes revealed pathways involved in keratinization, keratinocyte differentiation, skin development and barrier formation in *Dsg1*^−/−^ skin (Figure 2E, F). Changes in genes involved in keratinocyte differentiation were validated by qRT-PCR (Supplemental Figure 3D). The enrichment analysis also revealed increased ErbB signaling, in accordance with previously observed upregulation in human Dsg1-deficient epidermal equivalents (19). *Krt6*, known to be up-regulated in inflammatory conditions and considered a critical barrier alarmin (27), was also significantly increased, as were small proline rich protein (*Sprr*) genes in accordance with the decrease in loricrin protein expression (28).

To complement the RNA-Seq and protein analyses indicating a potential barrier defect in *Dsg1*^−/−^ animals, we carried out barrier function assays. An increase in toluidine blue dye penetration, a measure of the outside-in barrier, was observed in *Dsg1*^−/−^ animals (Figure 2G). We also observed an increase in transepidermal water loss, a measure of inside-out barrier, in P1 *Dsg1*^−/−^ animals (Figure 2H). We next analyzed changes in cell shape in the stratum granulosum (SG)-1 layer in the E18.5 epidermis by staining skin whole mounts for F-actin using phalloidin, which labels cells in the SG1 brightly. Circularity of cells in the SG1 layer was significantly reduced in *Dsg1*^−/−^ animals compared to *Dsg1^+/+^* animals (Figure 2I-J). This observation is consistent with our previous findings of changes in cell shape in Dsg1-deficient 3D epidermal cultures and suggests irregular packing of cells in this layer (19).

### E18.5 *Dsg1*^−/−^ mouse and human SAM syndrome skin share Th17 skewed inflammatory signatures

In addition to genes involved in skin differentiation and barrier function, pathway analysis of upregulated genes in *Dsg1*^−/−^ animals revealed genes involved in inflammatory processes including neutrophil chemotaxis, IL-17 signaling and antimicrobial humoral response (Figure 3A). Comparison of the *Dsg1*^−/−^ transcriptome to the transcriptome from cytokine-treated human keratinocytes revealed similarities to IL-17A, IL-36A and IL-36G-stimulated conditions, while there was no overlap with IL-13 or IL-4-treated keratinocytes (Figure 3B). Genes present in these pathways were validated by qRT-PCR for *Il1β, Cxcl1, Cxcl2, S100a8* and *S100a9* (Figure 3C). The fact that these responses were observed in embryonic skin raises the possibility that loss of Dsg1, in the absence of an external stimulus, primes a pro-inflammatory program in keratinocytes.

**Figure 3.**
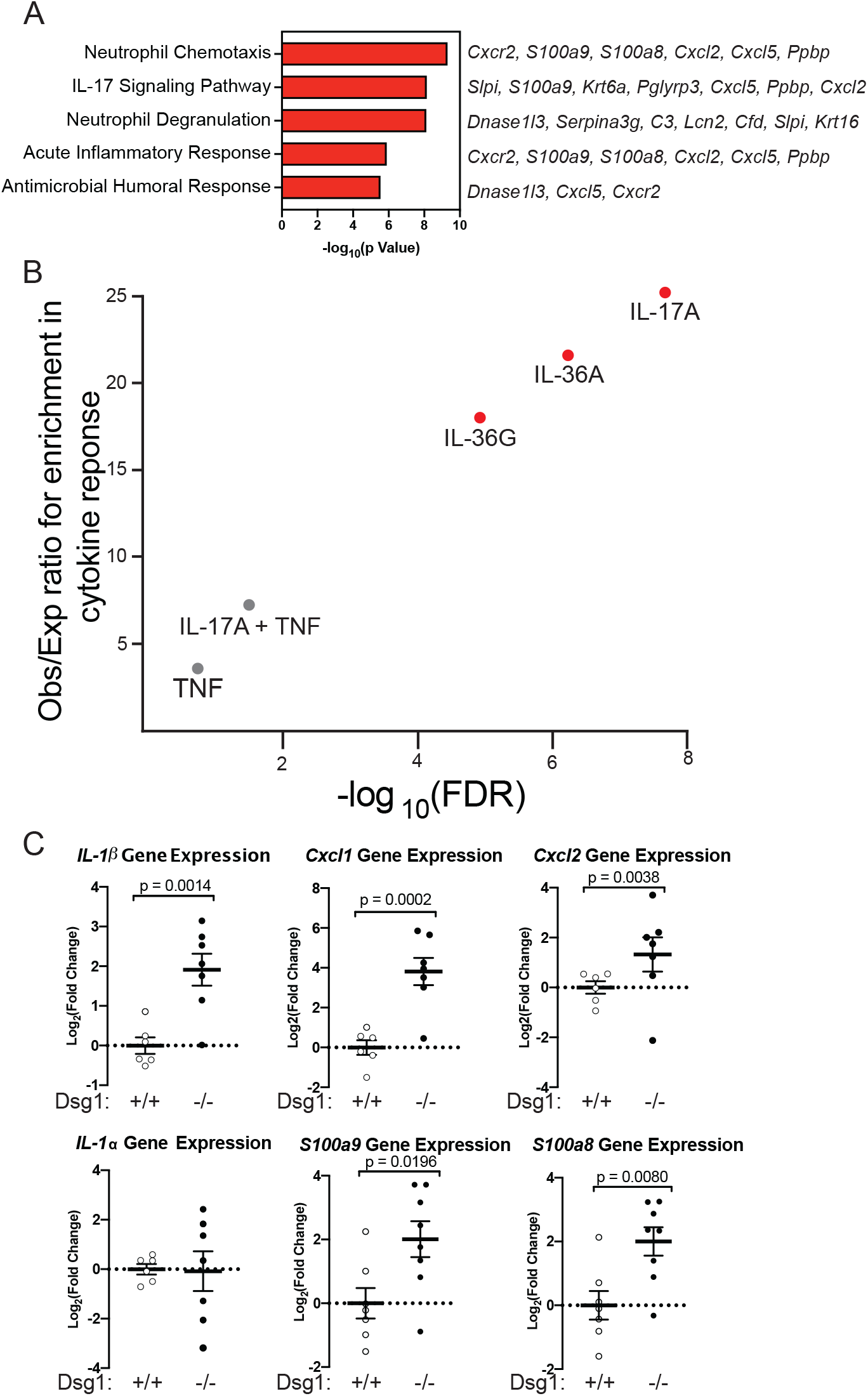
Dsg1 loss is associated with increased inflammatory response and anti-microbial pathways in E18.5 mouse epidermis. A) GO Biological Process terms significantly overrepresented in upregulated genes involved in immune processes from *Dsg1*^−/−^ skin. Gene names to the right of bars represent genes upregulated in *Dsg1*^−/−^ skin that belong to the GO associated pathway. B) Comparison between cytokine stimulated keratinocytes with *Dsg1*^−/−^ mouse skin. C) qRT-PCR for inflammatory cytokines and chemokines in mRNA from *Dsg1^+/+^* and *Dsg1*^−/−^ mouse skin.

To address the role of Dsg1 in humans, we took advantage of SAM syndrome patients with homozygous Dsg1 loss of function mutations. We performed whole transcriptome analysis on lesional skin biopsies taken from 4 patients, with 3 different Dsg1 mutations that result in the improper expression and/or delivery of the protein to the cell-cell membrane. These mutations include a missense mutation causing a premature stop codon, truncating part of Dsg1 at amino acid 887, a frame shift mutation, resulting in a truncation of the Dsg1 amino acid 621, and two patients with mutations leading to exon 2 skipping, which removes the Dsg1 signal sequence and leads to Dsg1 mislocalization (Supplemental Figure 4) (10, 12, 29). Principle component analysis (PCA) revealed that the four SAM syndrome patient lesional samples clustered away from the four control samples (Figure 4A). We also performed RNA-Seq on one non-lesional sample paired with one of the patient samples. PCA showed that this non-lesional sample was separated from both normal control samples and lesional samples. 767 upregulated and 1,701 downregulated genes (FDR ≤10% and |log_2_ Fold Change (FC)| ≥ 1) were present in lesional skin compared to control samples (Figure 4B).

**Figure 4.**
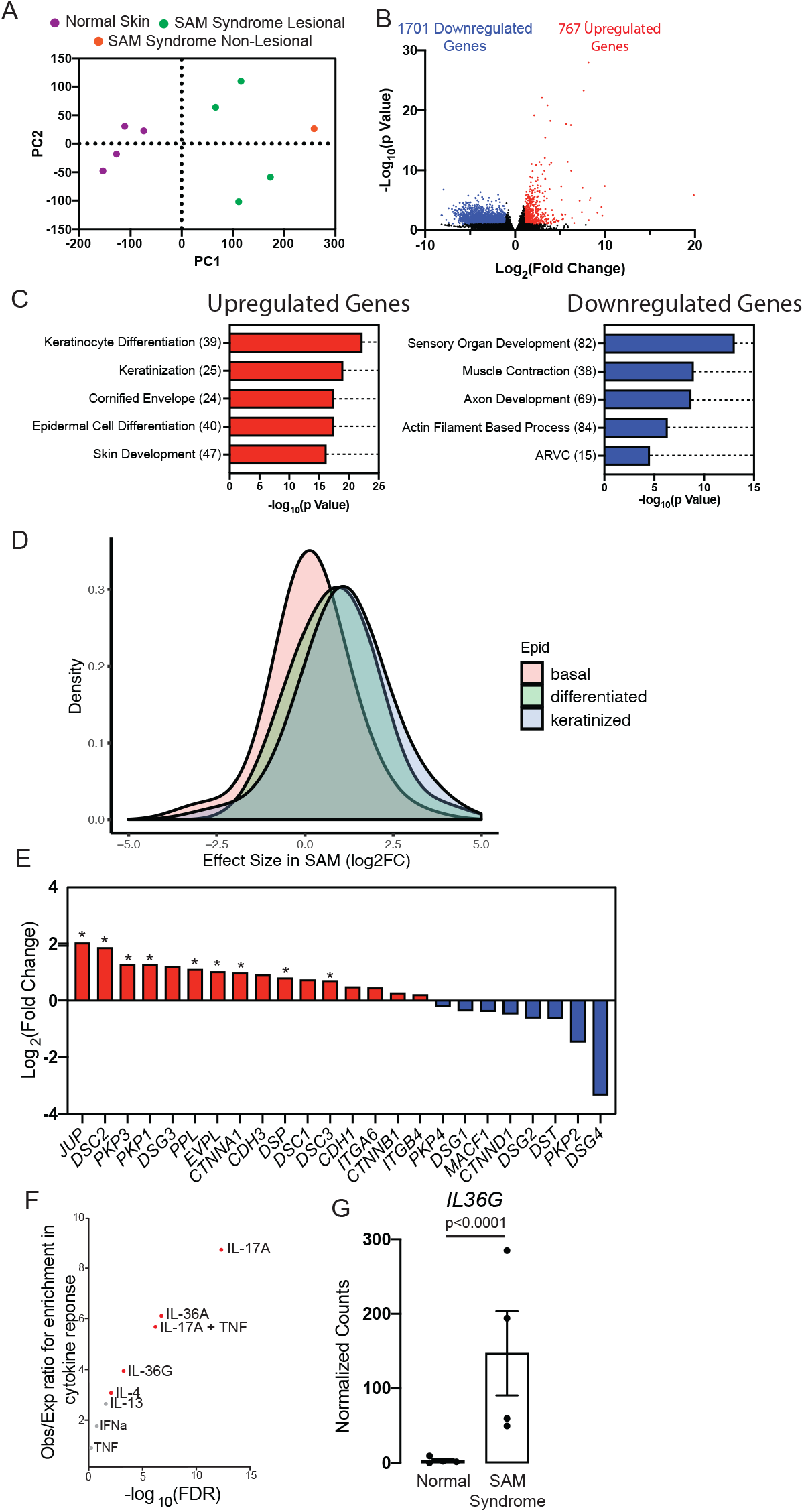
SAM syndrome patient skin whole transcriptome profile shows similarities to that of the *Dsg1*^−/−^ mouse. A) Principle component analysis (PCA) for SAM syndrome patient samples, 4 from lesional skin, 1 from non-lesional skin, and 4 normal skin whole transcriptome profiles. B) Volcano plot of upregulated and downregulated genes in RNA-Seq data from SAM lesional skin biopsy sections. Genes are considered significantly changed if they are greater than two-fold increased or decreased and have p < 0.1. C) GO Biological Process terms significantly overrepresented in upregulated (Red) or downregulated (Blue) genes from SAM patients. Value in parentheses represents the number of genes associated with each pathway. D) SAM syndrome RNA-Seq data was compared to a single cell RNA-Seq dataset from skin. The level of expression for each gene was determined in normal keratinocytes at different stages of differentiation and the expression level of these genes in SAM syndrome was graphed. D) Gene expression for desmosomal, adherens junction and hemidesmosome genes in the SAM data set. E) Cytokine response gene sets in keratinocytes overlapping with up-regulated genes in SAM syndrome. F) *IL36G* gene expression in SAM syndrome from RNA-Seq dataset G) *IL36G* gene expression in SAM syndrome from RNA-Seq dataset

Similar to what was observed in the *Dsg1*^−/−^ mouse skin, pathway analysis on upregulated genes from SAM syndrome patients found pathways involved in skin development, differentiation, keratinization and cornified envelope formation (Figure 4C). Downregulated genes from SAM syndrome patients partitioned into categories for neuron development, including sensory organ development and axon development, as well as pathways associated with the actin cytoskeleton (Figure 4C). Comparison to a single cell RNA-Seq data set from skin revealed that genes associated with differentiated and keratinized keratinocytes are upregulated in the skin in SAM syndrome patients, potentially due to an increase in the representation of these cells or a general upregulation of these genes in the upper epidermal layers (Figure 4D). Also observed were increases in RNAs encoding for desmosomal associated proteins, including *JUP*, *DSC2*, *PKP3*, *PKP1*, and *DSC3* and a decrease in *DSG4* (Figure 4E).

Comparison of upregulated genes in SAM syndrome with genes upregulated by specific cytokines in keratinocytes revealed a significant similarity to IL-17A and IL-36G responses, with the latter upregulated in the RNAseq dataset as well (Figure 4F, G) (30, 31). While IL-17A signaling has been implicated in both allergic inflammation and psoriasis, we observed less enrichment of genes involved in the pro-allergic IL-4 and −13 responses, in spite of the fact that patients with SAM syndrome often develop food allergies and have elevated levels of IgE (10, 22). Like the changes in keratinocyte differentiation pathways, these IL-17 signaling responses are similar to those observed in the *Dsg1*^−/−^ mouse skin (Figure 3B).

### Pemphigus foliaceus patient skin lacks the inflammatory profile shared by *Dsg1*^−/−^ mice and SAM syndrome patients

Pemphigus foliaceus (PF) is an autoimmune disorder in which circulating anti-Dsg1 autoantibodies bind to Dsg1 and interfere with cell-cell adhesion through steric hindrance and/or by stimulating endocytic turnover (6). Autoantibody mediated interference of Dsg1 results in loss of epidermal adhesion, which appears similar to that caused by genetic loss of Dsg1. Therefore, PF offered an opportunity to address whether genetic and autoantibody-induced interference of Dsg1 adhesion result in similar transcriptome remodeling. To date, transcriptome and cytokine analyses have focused primarily on circulating lymphocytes rather than the skin of PF patients (32). Towards this end, we carried out RNA-Seq on samples from seven PF patients. Compared to control samples, there were 461 significantly downregulated and 344 significantly upregulated genes in PF patient samples (FDR ≤10% and |log_2_ FC| ≥ 1, Figure 5A). Pathways revealed by GO analysis of significantly upregulated genes included epidermis development, establishment of skin barrier and keratinization (Figure 5B). Pathways involved in immune responses, such as regulation of leukocyte differentiation, were also observed in PF patient samples (Figure 5B). However, these pathways were distinct from those seen in SAM syndrome or the *Dsg1*^−/−^ mouse (Figure 4F, Figure 3A). Pathway analysis of downregulated genes revealed an enrichment for actin- and muscle-related processes and actin filament-based movement (Figure 5C). Comparison to a single cell RNA-Seq dataset from skin revealed that genes expressed by keratinized keratinocytes were generally upregulated in PF skin, without an enrichment in genes representing other keratinocyte populations (Figure 5D). The increase in keratinized genes (late differentiation) is similar to that observed in SAM syndrome; however, SAM syndrome also had an increase in keratinocyte specific genes representing earlier differentiation stages that were not observed in PF (Figure 4D). Analysis of cadherin and cadherin-associated genes in PF patient samples revealed similarities to the *Dsg1*^−/−^ mouse, including a significant increase in *DSC1* gene expression, with differences including significant downregulation of *DSG3* in PF patients compared to its upregulation in the *Dsg1*^−/−^ mouse (Figure 5E, Figure 1M). When compared to genes upregulated by specific cytokines in keratinocytes there was a moderate but significant overlap with IL-17A-treated cells; however, the strength of the association was much weaker in PF than the association in SAM syndrome and the *Dsg1*^−/−^ mouse, and no association with IL-36G or IL-36A responses was observed (Figure 5F).

**Figure 5.**
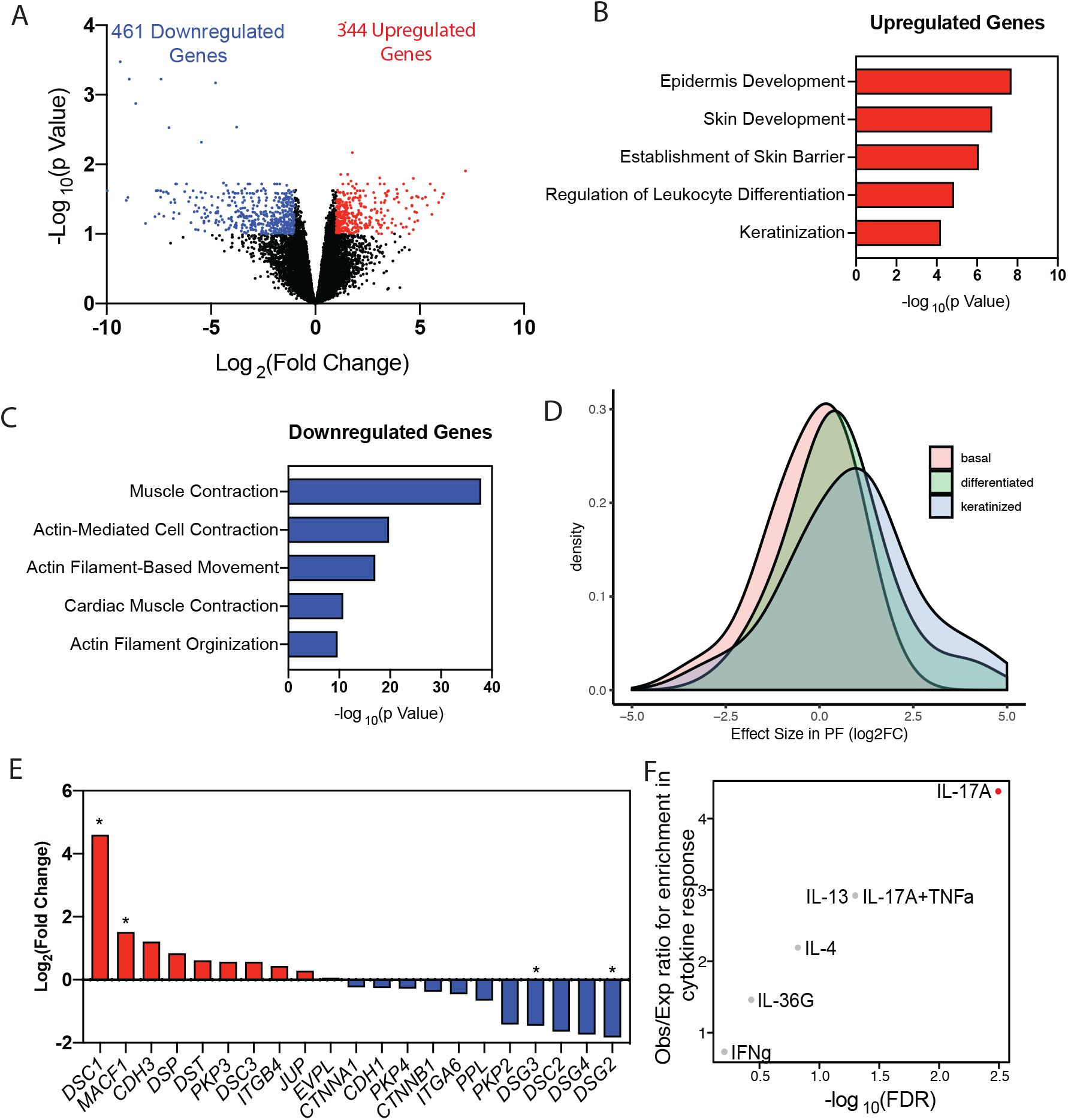
Whole transcriptional analysis of pemphigus foliaceus (PF) patient skin reveals profiles that are distinct from that of genetic Dsg1 deficiency. A) Volcano plot of upregulated and downregulated genes in RNA-Seq data from PF patient samples. B) GO Biological Process terms significantly overrepresented in upregulated genes from PF patients. C) GO Biological Process terms significantly overrepresented in downregulated genes from PF patients. D) PF patient RNA-Seq was compared to a single cell RNA-Seq data set from skin. The levels of expression for each gene was determined in normal keratinocytes at different stages of differentiation and the expression level of these genes in PF was graphed. E) Gene expression for desmosomal, adherens junction, and hemidesmosome genes in the PF data set. F) Cytokine response pathways in upregulated genes in PF patients.

Additional comparisons between the *Dsg1*^−/−^ mouse and SAM syndrome or PF revealed that ~25% of the genes significantly upregulated in *Dsg1*^−/−^ skin are also significantly upregulated in lesional skin from SAM syndrome, and ~15% of significantly downregulated genes in the *Dsg1*^−/−^ skin are also significantly downregulated in SAM syndrome (Figure 6A, B). Genes significantly upregulated in both SAM syndrome and *Dsg1*^−/−^ skin are genes expressed by differentiated keratinocytes (*SPRR* genes, *LCE3D*) or genes involved in antimicrobial responses in the skin (*S100A8*, *S100A9*, and the secretory leukocyte proteinase inhibitor of neutrophil elastase *SLPI*) (Supplementary Figure 5A). The overlap in expression patterns was present even though comparisons were between human skin from adults or adolescents and embryonic mouse skin, suggesting that Dsg1 regulates inflammatory pathways independently of an external stimulus, priming the skin for a proinflammatory response. Comparison between *Dsg1*^−/−^ skin and PF showed that only ~10% of genes significantly upregulated in the mouse are also significantly upregulated in PF, while only one gene was significantly downregulated in both conditions (Figure 6A, B, Supplementary Figure 5B). None of the inflammatory genes upregulated in the *Dsg1*^−/−^ mouse and SAM syndrome, such as *S100A8, S100A9*, and *SLPI*, were upregulated in PF patient skin. Thus, while all conditions are associated with changes in keratinocyte differentiation pathways, SAM syndrome and the *Dsg1*^−/−^ mouse share greater similarities in inflammatory signatures compared to PF.

**Figure 6.**
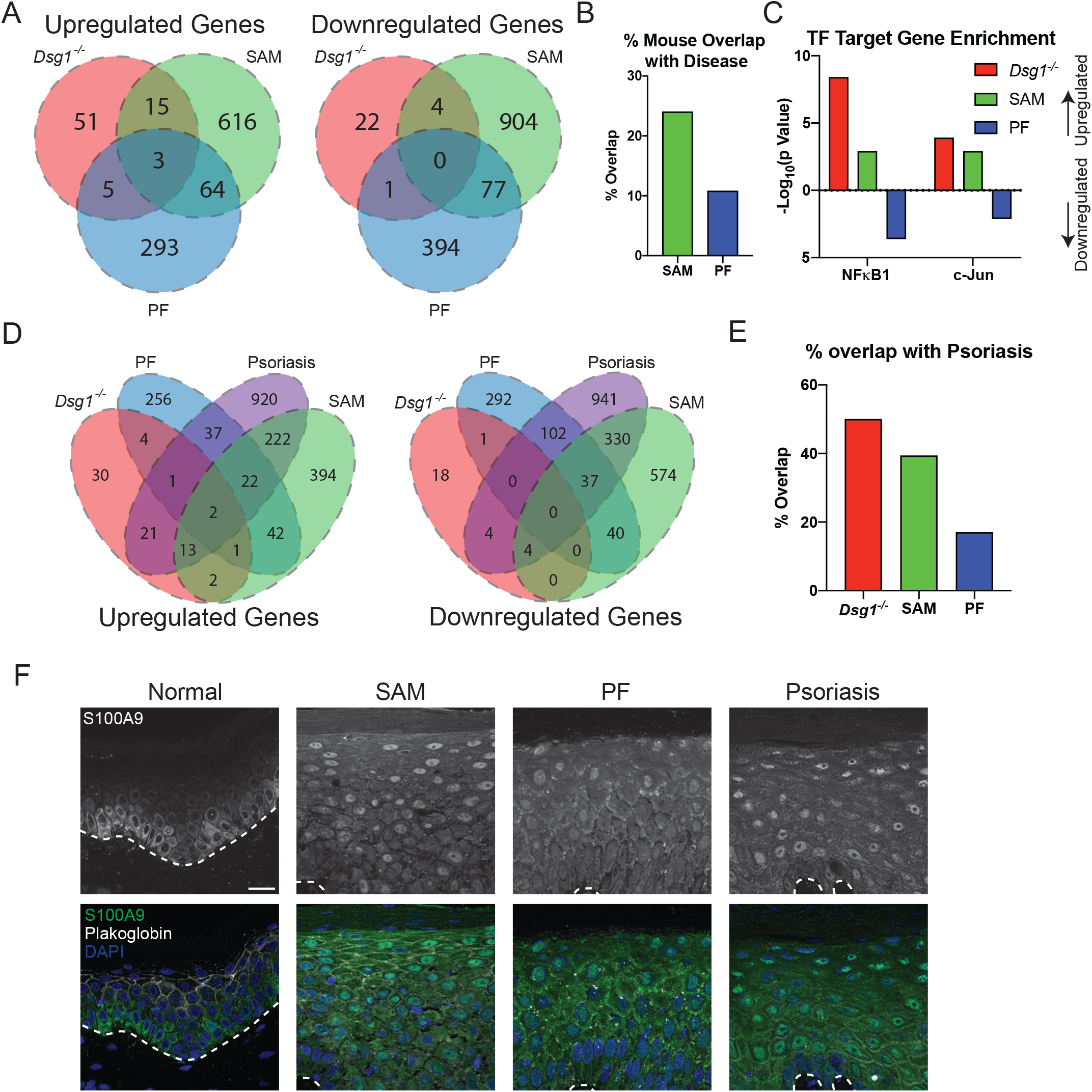
The inflammatory response profiles from *Dsg1*^−/−^ mice and SAM syndrome show similarity to psoriasis. A) Overlap between genes upregulated or downregulated in the *Dsg1*^−/−^ mouse skin, SAM syndrome and PF patients. B) Percent of genes upregulated in the *Dsg1*^−/−^ mouse skin and SAM syndrome or PF patients. C) Predicted transcription factor activity of NFκB1 and c-Jun. p values above the x axis represent enrichment of genes targeted by the transcription factor in the upregulated genes, while p values below the x axis represent enrichment of transcription factor targets in the downregulated genes. D) Overlap between genes upregulated or downregulated in *Dsg1*^−/−^ mouse skin, SAM syndrome, PF and psoriasis patients. E) Percent of genes upregulated in each disease that are also upregulated in psoriasis. F) Immunostaining for S100A9 and Plakoglobin to label cell membranes in control, SAM syndrome, PF and psoriasis patients. Scale bar = 20 μm.

We also analyzed predicted transcription factor activity on the upregulated and downregulated genes from all three data sets using the TRRUST database through Metascape. In both *Dsg1*^−/−^ mouse and SAM syndrome patients we observed a significant enrichment in upregulated genes regulated by NFκB1 and c-JUN (Figure 6C). This observation fits with the observed increase in inflammatory response pathways in these datasets. In contrast, we observed an enrichment in NFκB1 and c-JUN regulated genes in the downregulated genes in PF patients, consistent with the observation that the inflammatory response pathways are more weakly activated in PF patients (Figure 6C).

### Inflammatory profiles in *Dsg1*^−/−^ Skin and SAM syndrome, but not pemphigus foliaceus, are similar to psoriasis

Since inflammatory signatures exhibited by *Dsg1*^−/−^ animals and SAM syndrome patients are reminiscent of those observed in common inflammatory disorders such as psoriasis and atopic dermatitis, we compared the *Dsg1*^−/−^ transcriptome to those from 36 patients with psoriasis or atopic dermatitis. After pairing mouse genes with human orthologues, the top 12 patients with the strongest resemblance to *Dsg1*^−/−^ mice were from psoriasis (PSO) comparisons (Supplemental Figure 6A). Conversely, the 8 bottom-ranked with weakest resemblance (rs ≤ −0.00011) were from atopic dermatitis (AD) comparisons (Supplemental Figure 6A). Consistent with this, the 100 genes most strongly elevated in PSO lesions overlapped significantly with those genes elevated in *Dsg1*^−/−^ animal. This was not the case for genes elevated in AD lesions (Supplemental Figure 6B). In contrast, the 100 genes decreased in PSO or AD lesions did not overlap significantly with those genes altered in *Dsg1*^−/−^ mice (Supplemental Figure 6C). Direct comparison of *Dsg1*^−/−^ skin with psoriasis showed that ~50% the genes upregulated in *Dsg1*^−/−^ skin are also upregulated in psoriasis patient samples and that ~30% of genes downregulated in the *Dsg1*^−/−^ skin are also downregulated in psoriasis patients (Figure 6D, E). Genes upregulated in both psoriasis patient samples and *Dsg1*^−/−^ skin are involved with antimicrobial response (e.g. *S100A8, S100A9*, *SLPI*, *LCN2*), keratinization (e.g. *SPRR* genes), and leukocyte chemotaxis (e.g. *CXCL2, CXCL5, CXCR2*) (Supplementary Figure 6D). Comparison of whole transcriptome data from SAM syndrome patients with psoriasis patients showed a significant correlation between differentially expressed genes (Figure 6D, E, Supplemental Figure 6E, F). Direct comparison of significantly changed genes found that ~40% of genes upregulated in SAM syndrome are also upregulated in psoriasis, and ~35% of genes downregulated in SAM syndrome are downregulated in psoriasis (Figure 6D, E). In contrast, comparison of PF and psoriasis data revealed fewer overlapping genes (Figure 6D, E). When compared to cytokine response gene sets, we observed a significant enrichment in IL-17A, IL-36A and IL-36G response genes in psoriasis patients, as observed in SAM syndrome and the *Dsg1*^−/−^ mouse (Supplemental Figure 6G, Figure 3B, 4F).

These observations suggest that the skin in the *Dsg1*^−/−^ mouse and SAM syndrome share greater similarity with the inflammatory profile in psoriasis than that in PF patients. To validate these observations we stained skin samples from SAM syndrome, PF and psoriasis patients for S100A9. S100A9 is an antimicrobial peptide known to be elevated in psoriasis. *S100A9* mRNA was also increased in SAM syndrome and the *Dsg1*^−/−^ mouse, but not in PF patient samples (Supplemental Figure 5A). As previously described, S100A9 protein was increased in the skin of psoriasis patients compared to control, with high intensity nuclear staining present in the full thickness of the epidermis (Figure 6F)(33). High intensity nuclear S100A9 staining was also observed in SAM syndrome patient skin, but less so in PF patient skin (Figure 6F).

### Dsg1 downregulation is common in SAM syndrome and psoriasis

Based on the observation that the transcriptional profiles from *Dsg1*^−/−^ animals and SAM syndrome shared similarities with psoriasis we tested if Dsg1 levels were changed in psoriasis. Psoriasis samples from lesional skin exhibited a decrease in membrane associated Dsg1 staining compared to control or non-lesional skin from psoriasis patients, while Ecad levels were unchanged (Figure 7A, B, Supplemental Figure 6H-J). Previously we showed that Dsg1 regulates the stability of the gap junction protein, connexin 43 (Cx43). Consistent with this, membrane levels of Cx43 staining were decreased in lesional skin from psoriasis patients compared to control skin (Figure 7A, B). These observations raise the possibility that downregulation of Dsg1 in psoriasis has functional consequences for keratinocyte behavior. Reduction of Cx43 levels were also observed in PF patient epidermis, suggesting that our reported dependence of Cx43 stability on Dsg1 holds true across genetic and autoimmune disease (Figure 7C, D).

**Figure 7.**
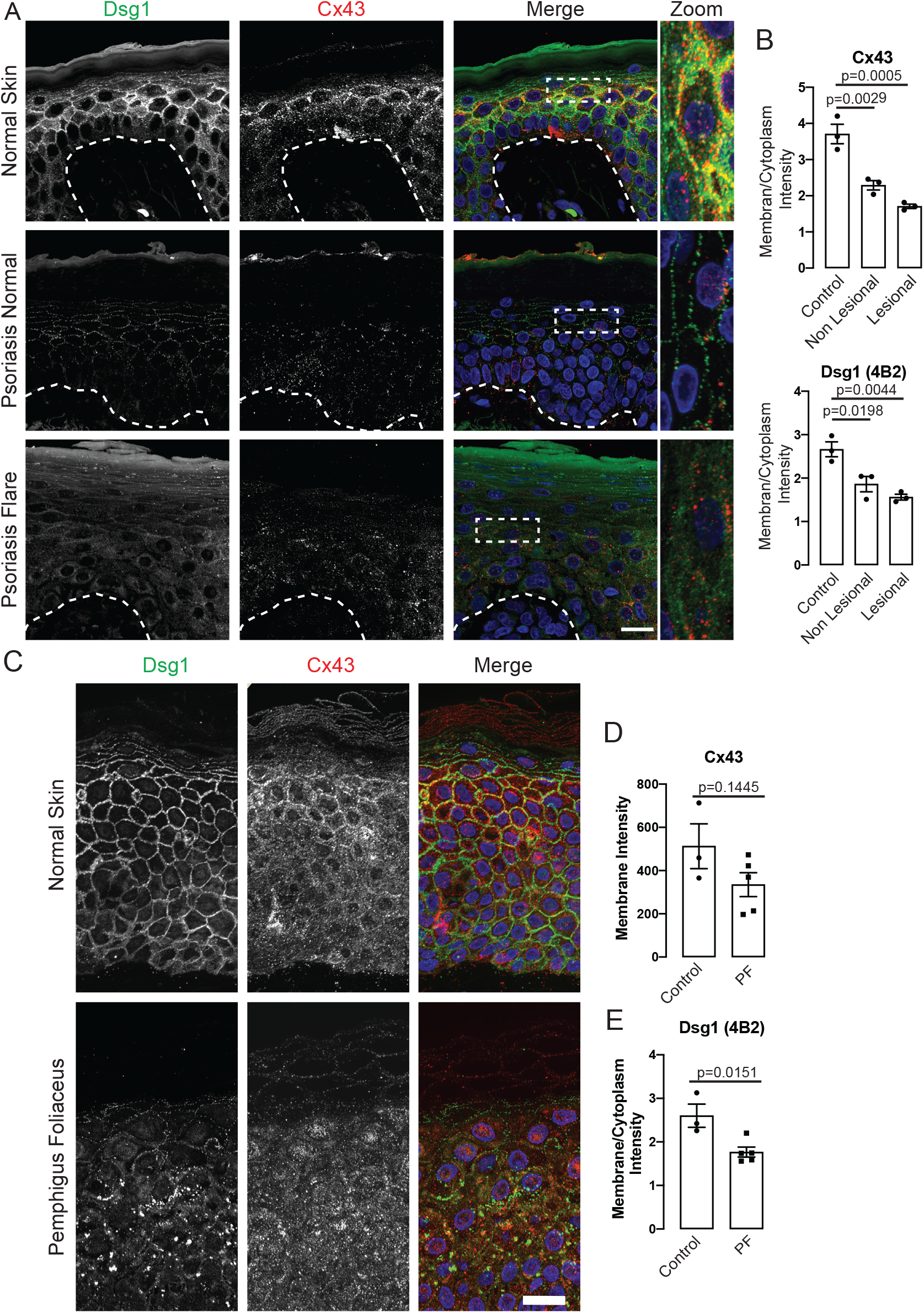
Loss of Dsg1 and Cx43 is common in psoriasis, SAM and pemphigus foliaceus. A) Immunostaining for Dsg1, using the cytoplasmic domain antibody 4B2 and Cx43 in normal skin, non-lesional skin and lesional skin from psoriasis patients. Scale bar = 20 μm. B) Quantification of Cx43 and Dsg1 immunostained samples expressed as membrane intensity over cytoplasmic intensity (n = 3). C) Immunostaining of Dsg1 using the 4B2 antibody, and Cx43 in normal skin and skin from PF patients. Scale bar = 20 μm. D) Quantification of Cx43 immunostained samples expressed as membrane intensity in PF and control samples (n=3-5). E) Quantification of Dsg1 immunostained samples expressed as membrane intensity over cytoplasmic intensity (n=3-5).

## Discussion

The epidermis is an immune organ that responds to environmental toxins, pathogens and mechanical stress through the coordinated activities of both keratinocytes and immune cells (34). Evidence is now emerging that cytoarchitectural components in keratinocytes are important contributors to keratinocyte responses to external stimuli. These architectural elements include desmosomes and their associated keratin intermediate filaments. For instance, proliferation-associated keratins K6, K16/17, which are turned on in response to a variety of stresses and in disorders such as psoriasis, have special structural roles but also act as alarmins that help stimulate innate immune defense and adaptive immunity (27). Intermediate filament anchoring desmosomes appeared during evolution around the time that adaptive immunity developed in jawless fish, and the keratinocyte specific desmosomal cadherin Dsg1 appeared even later when vertebrates became terrestrial and required a barrier suited to a new environment (18). We recently proposed the idea that this “newest” desmosomal cadherin acts as a sensor of environmental stress by remodeling the secretome upon stress-induced down-regulation of Dsg1 (16, 18). Supporting this, chronic loss of Dsg1, as occurs in SAM syndrome, is associated with chronic inflammatory and allergic disease. The extent to which keratinocyte Dsg1 directly controls these inflammatory and allergic responses and a systematic analysis of genes controlled by Dsg1 has not been explored in an *in vivo* setting.

Here we show that beyond its essential role in maintaining tissue integrity through desmosomal cell-cell adhesion in granular layers (35), Dsg1 controls the epidermal differentiation program and inflammatory gene expression. Importantly, the elevation of an inflammatory program occurs during embryogenesis, suggesting that Dsg1 may control these responses independently of any response to environmental stimuli.

Our previous work in an *in vitro* model of human epidermal morphogenesis demonstrated that in the absence of Dsg1 the epidermal differentiation program is impaired. This impairment is due to the loss of Dsg1-dependent attenuation of the EGFR/MAPK pathway and consequent failure of expression of superficial layer genes (19, 20). Global transcriptomics of skin from the Dsg1 deficient mouse described here showed up-regulation of ErbB and MAPK pathways, consistent with results from this human model. Also in alignment with results from the human model, protein levels and staining intensity for loricrin are also decreased in *Dsg1*^−/−^ animals, and cell shapes are irregular, likely contributing to the observed barrier defects. On the other hand, whereas the differentiation-dependent cadherin Dsc1 was reduced and Dsg3 unchanged in human Dsg1-deficient cultures, they are elevated in the *Dsg1*^−/−^ mouse. Similarly, Dsc1 is elevated in SAM syndrome caused by Dsg1 mutations (36). It seems likely that these are compensatory responses occurring through transcriptional or post-transcriptional mechanisms. Indeed, compensation has been shown to be common in the setting of a genetic mutation, but not in the setting of loss of function through knockdown (37). This is speculated to occur through loss of negative feedback loops, a response that appeared during evolution to make robust biological systems. As Dsg1 appeared late in evolution and is down regulated by a number of environmental stressors (18), it is possible that more ancient Dsgs might be upregulated under these circumstances.

SAM syndrome patients with Dsg1 loss of function mutations have severe allergies, and keratinocytes isolated from patients produce Th2 cytokines (10). Dsg1 deficiency also contributes to an allergic disorder called eosinophilic esophagitis (EOE), which exhibits Th2 skewing (38), and some SAM syndrome patients exhibit EOE as well (10). In addition, T cells isolated from patients with an endemic form of PF exhibit a Th2-skewed cytokine profile (39). In spite of these indicators of Th2 expression, the up-regulated genes in the whole transcriptome analysis of *Dsg1*^−/−^ animals and SAM syndrome patients both exhibited upregulation of Th17 associated pathways and a response to IL-17. An increase in the IL-36 pathway was also observed, which has been associated with multiple inflammatory disorders, including psoriasis where it promotes Th17 skewing through recruitment of Th17 cells, as well as a reduction in the keratinization program (23).

In contrast, we did not observe a strong IL-17/36 signature in the transcriptome from skin biopsies of PF patients. This observation is consistent with the recently reported weaker association of Th17 pathways with PF compared to the related blistering skin disease pemphigus vulgaris, caused by anti-Dsg3 antibodies (40). While the genetic and antibody-induced disorders share similarities in skin lesion morphology and signs that epidermal differentiation and keratinization programs are altered, the PF skin transcriptome lacks a strong inflammatory signature. The potential involvement of NFκB1- and c-Jun in expression of upregulated inflammatory genes in *Dsg1*^−/−^ mice and SAM syndrome, but not PF, is consistent with their well-known role as pro-inflammatory transcription factors.

The Th17 skewing due to Dsg1-deficiency is reminiscent of recent observations that ichthyosis patients with various underlying genetic basis, all having previously reported links to AD, showed robust Th17/IL-23 skewing (41, 42). A difference here, however, is that the Th17 inflammatory response is observed before birth, thus supporting a primary role for Dsg1 in the response. In this regard, it is interesting to note that pediatric AD populations share certain features of psoriasis, such as Th17 skewed inflammation, while adults from European and American populations skew more towards Th2 inflammation (22).

Our observations also raise the question of whether Dsg1 loss may be a common factor in psoriasis and other ichthyoses with Th17 skewing, as is the case, for instance, in Netherton syndrome, in which Dsg1 is degraded due to loss of LEKTI-1 (lympho-epithelial kazal type related inhibitor type 5) function (43). Indeed, Dsg1 reduction was observed in a cohort of psoriatic patient specimens, along with other changes we have previously linked with Dsg1 loss, such as Cx43 mis-localization, consistent with the possibility that Dsg1 loss may contribute to cytokine profiles in this disorder. While we do not yet know how Dsg1 is lost in psoriatic lesions, we do know that Dsg1 is particularly sensitive to stress-induced down regulation compared with other desmogleins and cadherin family members. Dsg1 expression is reduced in response to certain cytokines such as IL-13 in EOE, where Dsg1 loss has been implicated in contributing to disease pathogenesis (38). Dsg1 loss also occurs in response to bacterial exposure (44) and UV irradiation (17). Dsg1 is a substrate for MMPs and ADAM family proteases, the latter of which are known to contribute to psoriatic phenotypes through activating EGFR and TNF (45). The observations that Dsg1 loss in utero is sufficient to stimulate a Th17 skewed inflammatory response and Dsg1’s unique sensitivity to loss via extrinsic stressors, suggest that Dsg1 loss may activate a protective response. This is consistent with our recent findings that UV exposure selectively downregulates Dsg1, and conditioned media from Dsg1-deficient keratinocytes stimulates increased melanin secretion in primary melanocytes, a protective response to UV exposure (16, 17).

This work may explain recent reports that three patients with SAM or SAM-like syndrome responded well to the IL-23 inhibitor ustekinumab or secukinumab, supporting the efficacy of targeting the Th17 response in these disorders(46, 47). Future work that continues to analyze shared and distinct features of Dsg1 deficiency and common skin disorders could provide an opportunity to develop more targeted therapeutic approaches for both rare and common inflammatory disorders. In addition, our work raises the possibility that Dsg1 reduction could be a biomarker for Th17 skewing and taken into consideration when designing therapeutic protocols for skin disorders.

## Materials and Methods

### Generation of *Dsg1a* and *Dsg1b* exon 2 deleted mouse models

For details regarding the generation of *Dsg1a* and *Dsg1b* exon 2 deleted mice see supplemental methods.

### Generation of *Dsg1a-c* null mice

Mouse lines harboring the deletion of the ~170 Kb, Desmoglein 1 tandem gene cluster (*Dsg1 c, a, b*) on chromosome 18 of the mouse were generated in the Northwestern University Transgenic and Targeted Mutagenesis Laboratory. Gene editing using CRISPR/Cas9 technology was used to generate the deletion. mRNA Cas9 (GeneArt, Invitrogen A29378) and IVT sgRNAs (*Dsg1c*-L2 TAAATGACCCGGGGATTAGT and *Dsg1b*-R2 GGTTCAGGGAGGCTTCCCGC) were injected at a concentration of 25, 12.5, 12.5ng/μl, respectively, into the cytoplasm of single-cell fertilized C57BL/6J zygotes to introduce double strand breaks 5’ (*Dsg1g*-L2-5’) and 3’ (*Dsg1b*-R2) of the genes of the Dsg1 tandem. Microinjected zygotes were transferred into recipient females, and the progeny analyzed for gene editing. One founder with the ~170 Kb desired deletion was identified, sequence verified, and backcrossed for 10 generations with no change in the phenotypes described here.

### Immunofluorescence and image acquisition

See supplemental methods for a detailed description of immunofluorescence and image acquisition.

### Electron Microscopy

See supplemental methods for a detailed description of electron microscopy methods.

### Barrier Function Assays

See supplemental methods for a detailed description of barrier function assays.

### RNAscope

See supplemental methods for a detailed description of RNAscope methods.

### RNA analysis of mouse tissues

See supplemental methods for a detailed description of RNA analysis of mouse tissues and supplemental table 1 for sequences of primers used.

### Whole Mount Immunostaining and Analysis

See supplemental methods for a detailed description of whole mount staining and analysis methods.

### Immunoblot analysis of proteins

See supplemental methods for a detailed description of immunoblot methods.

### RNA-Seq Expression Profiling of *Dsg1*^−/−^ mice

RNA for RNA-Seq expression profiling was collected from flash frozen dorsal skin from E18.5 mice using the Quick-RNA miniprep kit (Zymo Research) following manual homogenization using the Tissue Squisher (Zymo Research) in lysis buffer. cDNA was synthesized using 1 μg of RNA using the Superscript III First Strand Synthesis Kit (Life Technologies/Thermo Fisher). RNA quality was determined using the 2100 Bioanalyzer (Agilent). 1μg RNA from each sample was used for mRNA enrichment using NEBNext Poly(A) mRNA magnetic isolation module (E7490S, NEB). RNA-seq libraries were constructed using the purified mRNA samples with NEBNext Ultra II Directional RNA library Prep Kit for Illumina (E7760S, NEB). The quality of these RNA-seq libraries were validated using 2100 Bioanalyzer (Agilent), and sequencing was performed by NUseq Core Facility using the IIlumina Hiseq4000 (1×50bp).

### RNA-Seq expression profiling of SAM syndrome and pemphigus foliaceus samples

RNA was isolated from 10μm sections of formalin-fixed paraffin embedded blocks from 4 lesional and 1 non-lesional SAM Syndrome skin, healthy control skin (n=4), patients with established pemphigus foliaceous (n=7), and PF matched controls (n=4). RNA was extracted using the E.N.Z.A. FFPE RNA Kit (Omega Bio-tek). Samples were prepared using the Lexogen 3’ QuantSeq mRNA-seq Library Prep Kit FWD (Lexogen) and sequenced on the Illumina NovsSeq 6000 system (Illumina).

### RNA-seq data processing and analysis

See supplemental methods for a detailed description of RNA-seq data processing and analysis

### Additional Gene Expression Datasets

For a detailed description of additional datasets used this study see supplemental methods.

### Antibodies

See supplemental methods for a detailed list of antibodies used in this study.

### Statistics

Unless otherwise stated, statistical analysis was performed using one-way ANOVA, with a Tukey correction for multiple comparisons, with all groups compared in each experiment. p < 0.05 was considered statistically significant in all experiments unless otherwise stated.

### Study Approval

All housing, care and use of animals was handled according to the Northwestern and CEINGE Institutional Animal Care and Use Committee (Protocol ID IS00001419 and Health Ministry protocol 888-2019-PR). *Dsg1^+/−^* mice on a C57BL/6 background were housed in a barrier facility in a temperature-controlled room with a 12hr light cycle and given ad libitum access to food and water. Unless otherwise described, all mouse experiments were performed on skin harvested from E18.5 mice. Mice were mated overnight and were separated the following morning to generate timed pregnancies. Embryos were harvested on day E18.5 and dorsal skin was collected from the embryo pelts.

Pemphigus foliaceus patients, psoriasis patients, SAM syndrome patients and healthy adults provided informed consent for skin biopsies. Tissues were anonymized for analysis and collected under IRB# HUM00087890 at the University of Michigan, and IRB# STU00009443 at Northwestern University and IRB-##0086-15 at Emek Medical Center.

## Author Contributions

KJG: Conceptualization, Writing - original draft preparation, writing - reviewing and editing, funding acquisition. LMG: Conceptualization, Methodology, Formal Analysis, Investigation, Writing - reviewing and editing, Visualization. QRC: Formal Analysis, Investigation, Writing - original draft preparation, Writing - review and editing, visualization. JLK: Investigation, Formal Analysis. LCT: Data Curation, Formal Analysis, Visualization. JAB: Investigation, Formal Analysis. GNF: Investigation, Formal Analysis. SML: Data Curation, Investigation. JK: Investigation. ALH: Investigation, Formal Analysis. HEB: Investigation. MH: Formal Analysis. SA: Investigation. PWH: Resources. JLJ: Investigation. GU: Investigation, Formal Analysis. LTD: Methodology. WRS: Formal Analysis. RA: Methodology. ES: Resources, Writing - review and editing. XB: Methodology, Funding acquisition. ECB: Resources, Investigation, Formal Analysis. CM: Conceptualization, Funding Acquisition. JEG: Resources, Writing - review and editing, funding acquisition.

## Acknowledgements

This work was supported by the National Institutes of Health/National Institute of Arthritis and Musculoskeletal and Skin Diseases R01 AR041836 and National Institutes of Health R37 AR43380 and National Cancer Institute R01 CA228196 to KJG, National Institute of Arthritis and Musculoskeletal and Skin Diseases R01 AR075015 to XB, and Telethon Foundation (Italy) GEP15096 to CM. QRC was supported by National Institutes of Health T32 Training Grant (T32 CA070085). MH was supported by the National Institutes of Health T32 Training Grant (T32 CA009560). Additional support was provided by the JL Mayberry endowment to KJG. We acknowledge support and materials from the Northwestern University Skin Biology and Diseases Resource-Based Center supported by P30AR075049, and the Center for Advanced Microscopy/Nikon Imaging Center. All histology services were provided by the Northwestern University Mouse Histology and Phenotyping Laboratory which is supported by NCI P30- CA060553 awarded to the Robert H Lurie Comprehensive Cancer Center. JEG, LCT, PWH are supported by NIAMS P30–AR075043. Gloria Urciuoli is a PhD student within the European School of Molecular Medicine (SEMM).

