## Supplemental Material for "Th17-skewed inflammation due to genetic deficiency of a cadherin stress sensor"

**Supplemental Materials**

**Supplemental Methods**

**Generation of *Dsg1a* and *Dsg1b* exon 2 deleted mouse model**

To mimic patient mutations which cause SAM syndrome, we deleted exon 2 from *Dsg1a* and *Dsg1b* in the mouse. CRISPR/Cas9 technology was used to generate exon 2 deletions in mouse *Dsg1a* and *Dsg1b* genes using sgRNA recognizing identical intronic sequences flanking exon 2 in *Dsg1a* and *Dsg1b*, but not in *Dsg1c*. Four sgRNA sequences were identified using Chopchop (http://chopchop.cbu.uib.no) and the best combination of sgRNA was selected by preliminary experiments in NIH3T3 cells, sgRNA2-CACCGAGACACATTACTTTGACATG, sgRNA4-CACCGGGAGGAGTAATTATGTCAGG. The selected sgRNA oligonucleotides were subcloned in the BbsI site into pSpCas9(BB)-2A-GFP (PX458, #48138, Addgene) or pSpCas9(BB)-2A-Puro (PX459, #62988 Addgene) vectors. Cloning was verified by Sanger sequencing using U6F1 primer (TACGATACAAGGCTGTTAGAGAG) that read-through the region of the BbsI site.

Mouse embryonic stem cells (mESC) E14Tg2A.4 were cultured in GMEM medium supplemented with 12% FBS (Hyclone CHA30070L), 2mM Glutamine, 1mM Sodium Pyruvate, 1x non-essential amino acids, 50 µM β-mercaptoethanol, 1000 U/mL ESGRO Leukemia inhibitory factor (Sigma). mESC cells were transfected with both vectors using Lipofectamine 2000 (Gibco). 10,000 cells were plated in 10cm$^2$ plates. After 30 hrs cells were selected with 3 µg/ml puromycin for 48 hrs. Cells surviving selection were analyzed for GFP expression. After 10 days, 171 clones were verified for the absence of Cas9 integration and homozygosity for exon 2 in both *Dsg1a* and *b.* One clone was used for blastocyst injections at the Institute of Genetics and Biophysics (IGB, Naples, Italy). Ten chimeras were born with skin erosions and did not survive.

Exon 2 deletion was determined by PCR on genomic DNA using the following specific oligonucleotides: *Dsg1a* forward: TGACCCTCAGTGCAATACAAAA, *Dsg1a* reverse: CTGGGCATGTTGAATCCTGTAA, *Dsg1b* forward: TGAACACCCATTACATGCTTCC, *Dsg1b* reverse: TTCGAATGAAGAGGTGCCTTTA.

**Immunofluorescence and Image Acquisition**

Dorsal mouse skin, SAM syndrome patient, PF patient and healthy control skin samples were either fixed in 10% formalin overnight, embedded in paraffin blocks and cut into 4 µm thick sections, or embedded in Optimal Cutting Temperature compound (OCT, Tissue-Tek) and cut into 4-5 µm sections. For immunostaining, FFPE sections were baked at 60°C overnight and de-paraffinized using xylene. Samples were then rehydrated through a series of ethanol and PBS dips, and slides were permeabilized in 0.5% Triton X-100 in PBS. Antigen retrieval was performed by incubation in 0.01 M Citrate buffer at 95°C for 15 minutes. Sections were blocked in blocking buffer (1% BSA, 2% normal goat serum in PBS) for 60 minutes at 37°C. Samples were then incubated in primary antibody at 4°C overnight, followed by incubation in secondary antibodies for 1 hour at 37°C. OCT samples were allowed to warm to room temperature and were fixed either with 4% paraformaldehyde or 100% anhydrous methanol. Staining for OCT samples followed the same method as FFPE samples, excluding the antigen retrieval steps. Images were acquired using an AxioVison Z1 system (Carl Zeiss) with Apotome slide module, an AxioCam MRm digital camera, and a 40x (0.5 NA, Plan-Neofluar) objective. Image analysis was carried out using ImageJ software. For IHC, I-View DAB detection kit (Ventana, Roche, San Jose) was used according to the manufacturer's instructions.

For exon 2 deletion animals, immunohistochemistry was performed with the R.T.U. VECTASTAIN Universal Elite ABC Kit (VECTOR LABORATORIES PK-7200), following the manufacturer's instructions. Detection was performed with DAB Peroxidase Substrate (VECTOR LABORATORIES SK-4100). Tissue was counterstained with hematoxylin (Hematoxylin QS; VECTOR H-3404). Images were acquired using an Axioskop 2 Plus (Carl Zeiss) and AxioCam color digital camera 20x (0.5 NA, Plan-Neofluar) objective.

**RNA analysis of mouse tissues**

Mouse C57BL/6 organs were dissected and total RNA was extracted using TRIzol reagent (Invitrogen). cDNA was synthesized using SuperScript Vilo (Invitrogen). qRT-PCR was performed using the SYBR Green PCR master mix in an ABI PRISM 7500 (Applied Biosystems). Levels of the target genes were quantified using specific oligonucleotide primers and normalized for ß-actin.

RNA for qRT-PCR experiments was collected from flash frozen dorsal skin of E18.5 mice. RNA was isolated from flash frozen dorsal skin using the Quick-RNA miniprep (Zymo Research) following manual homogenization using the Tissue Squisher (Zymo Research) in lysis buffer. cDNA was synthesized using 1 μg of RNA using the Superscript III First Strand Synthesis Kit (Life Technologies/Thermo Fisher). Quantitative PCR was performed on the QuantStudio 3 instrument (Thermo Fisher), using SYBR Green PCR master mix (Thermo Fisher). Relative mRNA levels were calculated using the ΔΔCt method normalized to GAPDH. Primer sequences in Supplemental Table 1.

**Antibodies**

Antibodies used in this study include: mouse anti-Dsg1(P124, 651111, Progen; 27B2, 32-6000, Thermo Fisher; 4B2), mouse anti-Dsg3 ( D219-3, MBL international Corp.), anti-Dsc1 (U100, 61092, Progen), mouse anti-Ecad (610181, BD Biosciences (for mouse samples), mouse anti-Ecad (HECD1, for human samples, gift from M. Takeichi and O. Abe, Riken Center for Developmental Biology, Kobe, Japan), anti-Cx43 (AB1728, EMD Millipore), rabbit anti-transglutaminase (sc-25786, Santa Cruz Biotechnology), IL-23p19 (511201, Biolegend, USA), mouse anti-S100A9 (ab105472 Abcam), mouse anti-desmoplakin (91121236-1VL, Sigma Aldrich), chicken anti-plakoglobin (Aves Laboratories, Tigard, OR). The following antibodies were a gift from J. Segre (National Human Genome Research Institute, National Institutes of Health): rabbit anti-loricrin, rabbit anti-involucrin.

**RNAscope**

RNA *in situ* hybridization was completed using RNAscope technology with the ACDBio RNAscope 2.5HD Reagent Kit - Brown (ACDBio, Hayward, CA). FFPE tissues were sectioned at 4μm and hybridized with probes against *Dsg1a* (ACDBio, 84861) or *Dsg3* (ACDBio, 464301). The protocol was followed according to manufacturer's instructions and hybridized RNA was detected with DAB and counterstained with Gill's hematoxylin.

**Immunoblot analysis of proteins**

Immunoblot for Exon 2 knockout mice were performed as follows. Skin samples were lysed with a lysis buffer (6% SDS, 0.125M Tris-HCl pH 6.8, 1x Protease Inhibitor, 1x Phosphatase Inhibitor and 1x PMSF) supplemented with 20% β-mercaptoethanol. Samples were separated by SDS-PAGE. Transferred blots were then incubated with the Dsg1 B-11 antibody (Santa Cruz Biotechnology, sc-137164), anti-β-actin antibody for 2 hours at room temperature, then incubated with HRP conjugated secondary for 1 hour at room temperature.

All other immunoblots were performed as follows. Lysates were collected from E18.5 dorsal skin by manual homogenization using the Tissue Squisher (Zymo Research) in urea sample buffer (8M urea, 1% SDS, 60mM Tris (pH 6.8), 5% β-mercaptoethanol, 10% glycerol), sonicated and centrifuged. Samples were separated by SDS-PAGE and transferred to nitrocellulose membranes. Membranes were blocked with 5% milk in PBS, incubated with primary antibody overnight at 4°C, and secondary antibody conjugated to HRP for 1 hour at room temperature. Proteins were imaged using chemiluminescence on Odyssey FC imaging system (Licor). Densitometry values were analyzed using ImageStudio software (Licor) and normalized to GAPDH or Actin.

**Whole mount Imaging and Analysis**

Dorsal skin was harvested from E18.5 mice, and either fixed immediately for 2 hours in 4% paraformaldehyde or incubated with 2.4 U/mL dispase in PBS at 37°C for 1 hour. The epidermis was peeled from the dermis and fixed for 15 minutes with 4% paraformaldehyde. Samples were blocked in blocking solution (5% normal goat serum, 1% Triton X-100, in PBS) overnight at 37°C. Samples were then incubated with primary antibodies overnight at 37°C followed by incubation with Alexa-Fluor conjugated secondary antibodies and phalloidin-647 diluted in blocking solution overnight at 37°C. Samples were mounted onto glass slides with Prolong gold (Life Technologies). Z-stack images (z-step size of 1.5 μm) were taken on a Nikon A1R confocal laser scanning microscope with two PMT detectors and two GaAsP detectors using a 40x (1.0 NA, Oil) objective, controlled by NIS Elements software (Nikon). Analysis of cell circularity was performed using ImageJ software on phalloidin images.

**Additional Gene Expression Datasets**

The fold change signature of *Dsg1*$^{-/-}$ vs. *Dsg1*$^{+/+}$ skin was compared to 36 others generated from microarray experiments comparing PSO or AD lesions to normal or uninvolved human skin (Supplemental Figure 6A). Genome-wide fold-change estimates were calculated from each PSO/AD vs. normal/uninvolved comparison as described previously (*55*). Mouse genes were paired with their human orthologue based on the Homologene database (*56*), creating human-mouse orthologous gene pairs. Spearman's *rho* statistic was calculated for each comparison with p-values calculated based on the asymptotic t approximation. To identify genes robustly elevated by PSO and AD, we calculated meta-signatures by averaging fold-change estimates across the subset of comparisons that used the same Affymetrix Human Genome Plus 2.0 microarray platform (n = 11, PSO; n = 10, AD). Human genes without a mouse orthologue were excluded from this analysis. Based on the composite meta-signature, we identified the 100 genes most strongly increased by PSO and AD (i.e., highest average fold-change) and the 100 genes most strongly decreased by PSO and AD (i.e., lowest average fold-change). We then evaluated cumulative overlap between these 100 genes and the list of corresponding mouse genes ranked according to *Dsg1*$^{-/-}$ /*Dsg1*$^{+/+}$ fold-change.

**RNA-seq data processing and analysis**

Quality control and adaptor trimming were performed on sequence reads from the RNA-seq data. STAR alignment (*1*) was used to align reads to the reference (GRCh37 for human and mm10 for mouse samples). HTSeq (*2*) was used for gene quantification and DESeq2 (*3*) was used for normalization and differential expression analysis.

For the comparison between the different datasets (e.g. *Dsg1*$^{-/-}$ vs SAM), only genes that are common were considered in the analysis. To identify the cytokine signature of each skin condition, we took the genes induced by cytokines in human keratinocytes, and computed the enrichment among the top 500 most significant genes up-regulated in the corresponding skin condition using the hypergeometric test. Functional enrichment analysis was performed using Metascape (metascape.org) using the Gene Ontology (GO) pathways, and Kyoto Encyclopedia of Genes and Genomes (KEGG) pathways. Pathways were considered statistically significant with $p < 0.05$.

**Electron microscopy**

Dorsal skin from E18.5 embryos were processed for conventional electron microscopical analysis as described in (Scothern and Garrod, 2008). Briefly, dorsal skin was cut into pieces and fixed in 0.1M cacodylate buffer pH 7.3 containing 2% PFA and 2.5% glutaraldehyde overnight. Tissues were postfixed in 2% osmium tetraoxide followed by 2% uranyl acetate. Tissues were dehydrated in ascending grades of ethanol, infiltrated with propylene oxide and embedded in Embed 812 resin, cured overnight at 60°C. Tissues were ultrathin sectioned with Leica Ultracut UC6 ultramicrotome with a diamond knife and collected on formvar coated copper mesh grids. Sections on grids were stained in 3% uranyl acetate followed by Reynolds lead citrate solution. Stained sections were viewed and photographed using an FEI Tecnai Spirit G2 transmission electron microscope.

**Skin barrier toluidine blue assay**

For outside-in barrier testing, E18.5 embryos were sacrificed and rinsed in PBS followed by dehydration immersion steps of 25%, 50%, 75%, 100%, 75%, 50%, 25% methanol. After the dehydration steps the embryos were rehydrated in PBS and immersed in 1-5% toluidine blue for up to 10 minutes and washed several times with PBS.

**Transepidermal water loss (TEWL)**

Tewameter TM300 system (Courage + Khazaka electronic GmbH) fitted with a small animal adapter, was used to measure water evaporation from the skin surface and thus quantifying epidermal permeability barrier function of P1 pups over 5 hours on the dorsal back. Measurements were recorded when TEWL readings were stabilized at approximately 45 seconds after the probe was placed on the skin and readings were averaged from 5 readings per time point.

| Gene | 5' | 3' |
|---|---|---|
| ***Dsg1a*** | CAAGGCACTTCTTCCACTGAGA | CGCTGCCTCCCCATGA |
| ***Dsg1b*** | GGAGGCAGTGGAGTTAACAACAC | CGGGTTCCGGTTCGTCTA |
| ***Dsg1c*** | GACCTGCACAGGGACAATCA | GATGTTGTGGGATGTTCAGTAGTTG |
| ***Dsg3*** | CCTGACAGTGTGTCAATGTG | GGCTGAGCTCCTTCGATTCC |
| ***Dsg4*** | GCGGGGATTGATCGGCCACC | CTTGATTCTGCAGTCACATTC |
| ***Dsc1*** | GCTCTGCATTGCTACTGTGC | ACACCTTTTCACCAAGCCGA |
| ***Dsc3*** | ATGGTGGTTCCTGAGTTCCG | TTGAGGCGTGTGTGCATAGT |
| ***Pkp1*** | GCATAACCTCTCCTACCGCC | CCATTGGACATCAGCCCCTT |
| ***Pkp2*** | ACGAAGATGTTCAACGGGCT | CCGAGGCACTCCATTCAGTT |
| ***Pkp3*** | GCAAGCCTGAGACTGGTGTT | TCGCTCATGGAAGGACACTG |
| ***Dp*** | AGCTCGATGGAAAGTCAGCC | GGGAGAGTCTGTCCATCTGGT |
| ***Jup*** | GCTGCCCAGAGTATGATCCC | GGGGTAGTCTCCATCCAGGT |
| ***Lor*** | CTCCTGTGGGTTGTGGAAAGA | TGGAACCACCTCCATAGGAAC |
| ***Il1a*** | GCACCTTACACCTACCAGAGT | AAACTTCTGCCTGACGAGCTT |
| ***Il1b*** | GAAATGCCACCTTTTGACAGTG | TGGATGCTCTCATCAGGACAG |
| ***Cxcl1*** | ACTGCACCCAAACCGAAGTC | TGGGGACACCTTTTAGCATCTT |
| ***Cxcl2*** | CCAACCACCAGGCTACAGG | GCGTCACACTCAAGCTCTG |
| ***s100a8*** | CCTTTGTCAGCTCCGTCTTCA | TCCAGTTCAGACGGCATTGT |
| ***s100a9*** | GCACAGTTGGCAACCTTTATG | TGATTGTCCTGGTTTGTGTCC |
| ***Flg1*** | ATGTCCGCTCTCCTGGAAAG | TGGATTCTTCAAGACTGCCTGTA |
| ***Flg2*** | CTAGAGGGCATGAGTGTAGTCA | CAAGACTGGACAGTTGGCTGG |
| ***Ivl*** | ATGTCCCATCAACACACACTG | TGGAGTTGGTTGCTTTGCTTG |
| ***Tgm1*** | TCTGGGCTCGTTGTTGTGG | AACCAGCATTCCCTCTCGGA |
| ***Cdsn*** | TTGCTGATGGCCGGTCTTATT | GCCAGTCTTTCCAATGAGACAAG |
| ***Cdh1*** | CAGGTCTCCTCATGGCTTTGC | CTTCCGAAAAGAAGGCTGTCC |
| ***Cdnnb1*** | ATGGAGCCGGACAGAAAAGC | CTTGCCACTCAGGGAAGGA |
| ***Cdnna1*** | AAGTCTGGAGATTAGGACTCTGG | ACGGCCTCTCTTTTTATTAGACG |
| ***Cdh3*** | CTGGAGCCGAGCCAAGTTC | GGAGTGCATCGCATCCTTCC |
| ***Krt1*** | TGGGAGATTTTCAGGAGGAGG | GCCACACTCTTGGAGATGCTC |
| ***Krt10*** | CGAAGAGCTGGCCTACCTAAA | GGGCAGCGTTCATTTCCAC |
| ***Krt14*** | AGCGGCAAGAGTGAGATTTCT | CCTCCAGGTTATTCTCCAGGG |
| ***Krt5*** | TCTGCCATCACCCCATCTGT | CCTCCGCCAGAACTGTAGGA |

**Supplemental Table 1. Sequences for qPCR primers**

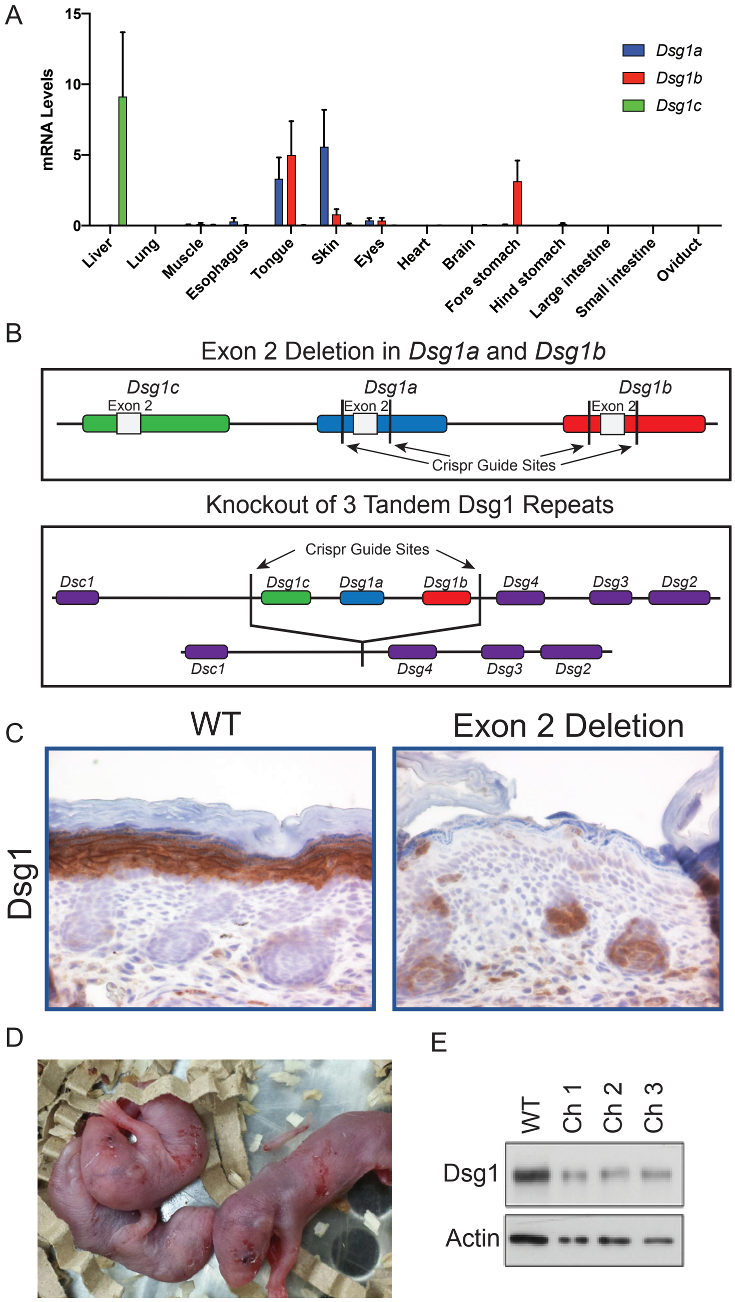


**Supplemental Figure 1. Exon 2 deletion in mice causes perinatal lethality and reduced Dsg1 in blistering areas.** A) Real-time qRT-PCR from RNA isolated from mouse organs demonstrating the expression for *Dsg1a*, *Dsg1b*, and *Dsg1c*. n = 3. B) Schematic representing the cadherin gene cluster in mice and describing the two knockout strategies pursued. C) Immunostaining for Dsg1 in the skin of *Dsg1*$^{+/+}$ and *Dsg1a* and *b* exon 2 deletion chimeras. D) Images of newborn chimeras demonstrating the presence of skin blisters. E) Immunoblot for Dsg1 in skin samples from 1 *Dsg1*$^{+/+}$ and 3 *Dsg1* exon 2 deletion chimeric animals. Actin was used as a loading control.

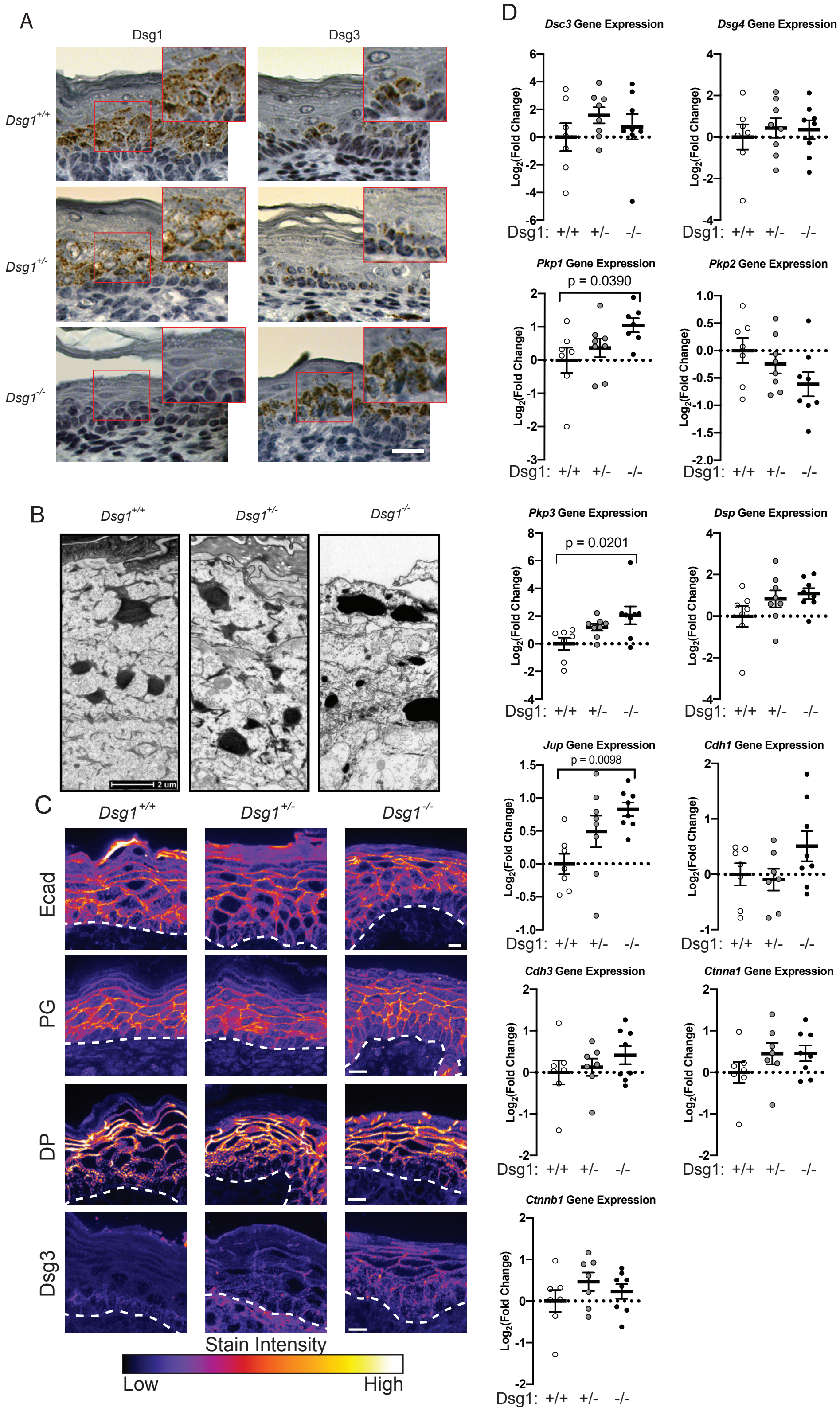


**Supplemental Figure 2. Dsg1 knockout perturbs normal keratinocyte adhesion and expression of adhesion molecules.** A) RNAscope showing *Dsg1* and *Dsg3* gene expression in E18.5 mouse skin. B) EM images of the skin from E18.5 mouse skin. C) Immunostaining for Ecad, plakoglobin (PG) and desmoplakin (DP) and Desmoglein 3 (Dsg3) in the skin from E18.5 mice. Scale bar = 20 μm. D) Gene expression for cadherin and cadherin associated genes. Fold change in gene expression was calculated using the ΔΔCT method, normalizing to GAPDH and then the $Dsg1^{+/+}$ mouse.

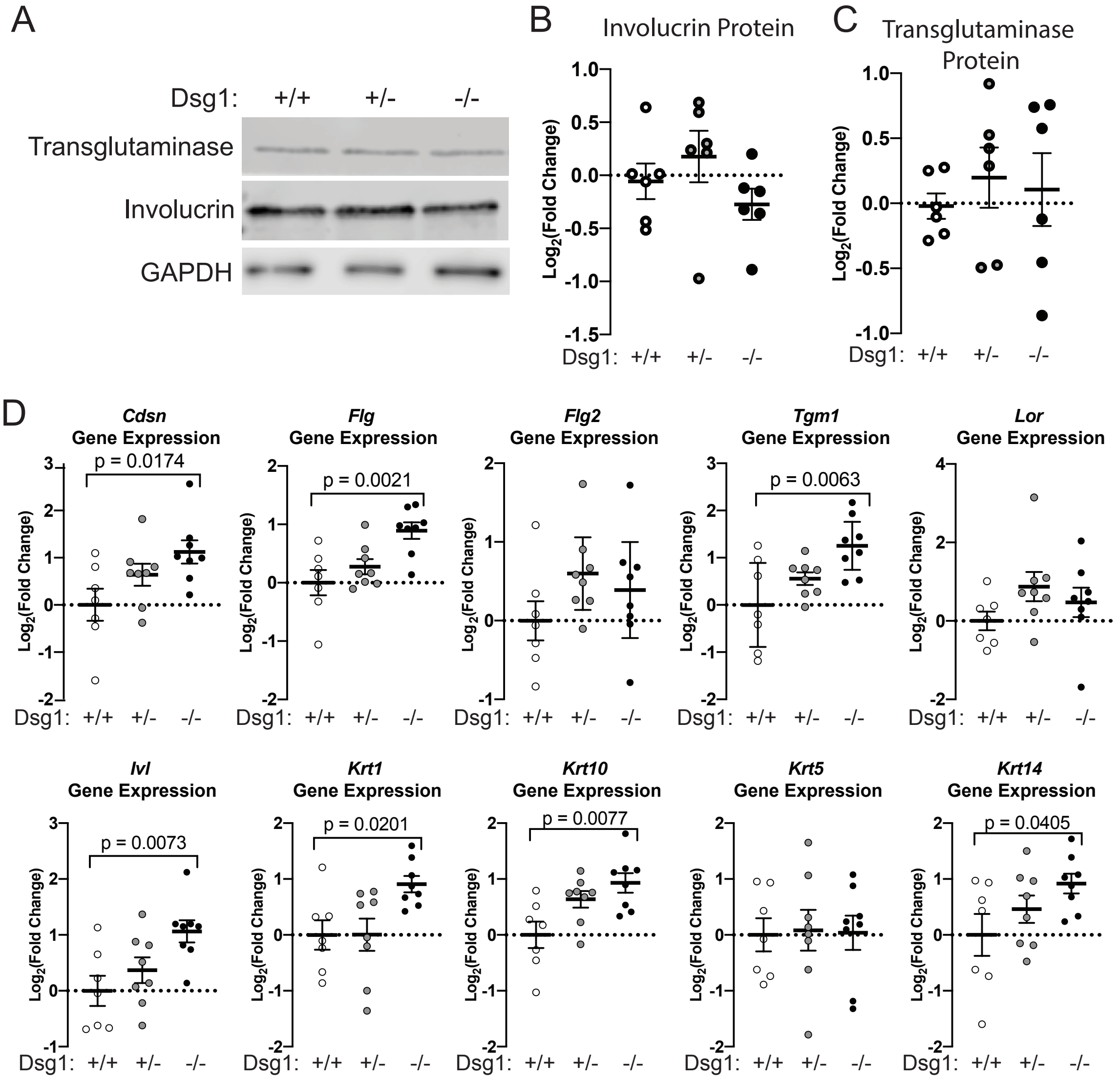


**Supplemental Figure 3. Dsg1 knockout disrupts normal keratinocyte differentiation.** A) Immunoblot for involucrin and transglutaminase in protein extracts from E18.5 mouse skin. GAPDH was used as a loading control. B) Quantification of involucrin protein from immunoblot. Densitometry values were normalized to the *Dsg1*$^{+/+}$ samples and GAPDH was used as a loading control (n = 6/genotype). C) Quantification of transglutaminase protein from immunoblot. Densitometry values were normalized to the *Dsg1*$^{+/+}$ samples and GAPDH was used as a loading control (n = 6/genotype). D) Gene expression for genes expressed in keratinocyte differentiation specific patterns. Fold change in gene expression was calculated using the ΔΔCT method, normalizing to GAPDH and then the *Dsg1*$^{+/+}$ mouse.

A

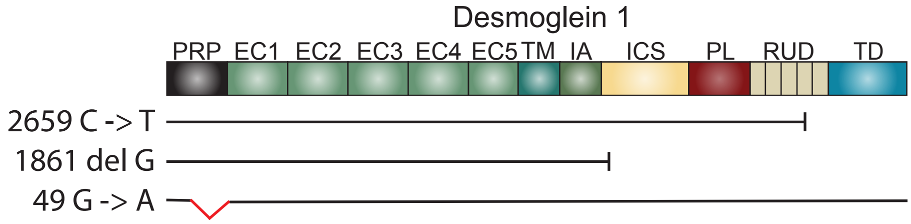


**Supplemental Figure 4. Dsg1 domains and associated mutations in SAM syndrome patients.** A) Schematic showing protein domains for Dsg1 and mutations associated with patients from SAM syndrome used in this manuscript. PRP, signal peptide and proprotein; EC, extracellular; EA, extracellular anchor; TM, transmembrane; IA, intracellular anchor; ICS, intracellular cadherin-like sequence; IPL, intracellular proline-rich linker; RUD, repeat unit domain; DTD, desmoglein specific terminal domain.

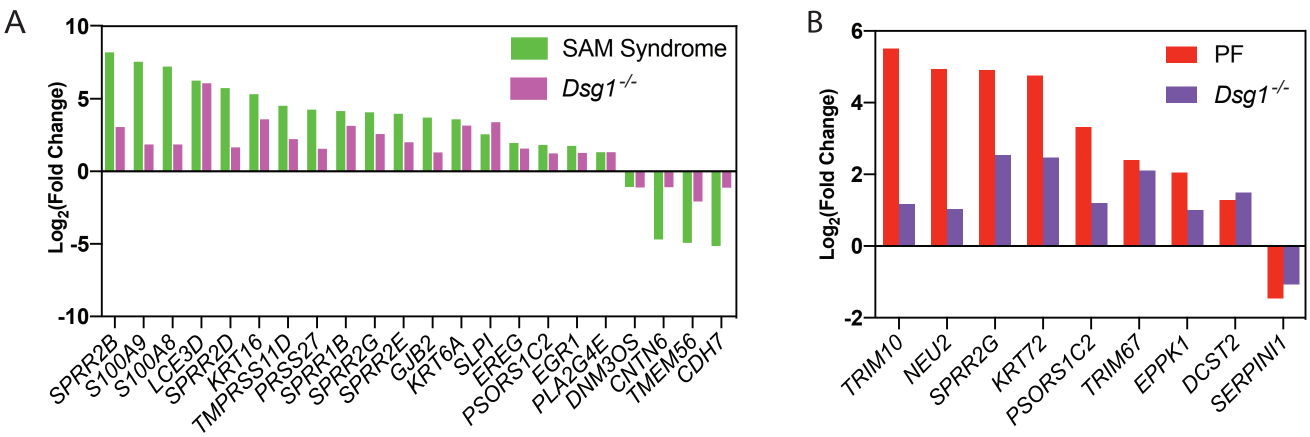


**Supplemental Figure 5. *Dsg1$^{-/-}$* mice are more similar to SAM syndrome than PF.** A) Fold change in genes significantly upregulated or downregulated in both SAM syndrome and in the *Dsg1$^{-/-}$* mouse skin. Genes were considered significantly changed if $|FC| > 2$, and $p < 0.1$. B) Fold change in genes significantly upregulated or downregulated in both *Dsg1$^{-/-}$* mouse skin and PF patients.

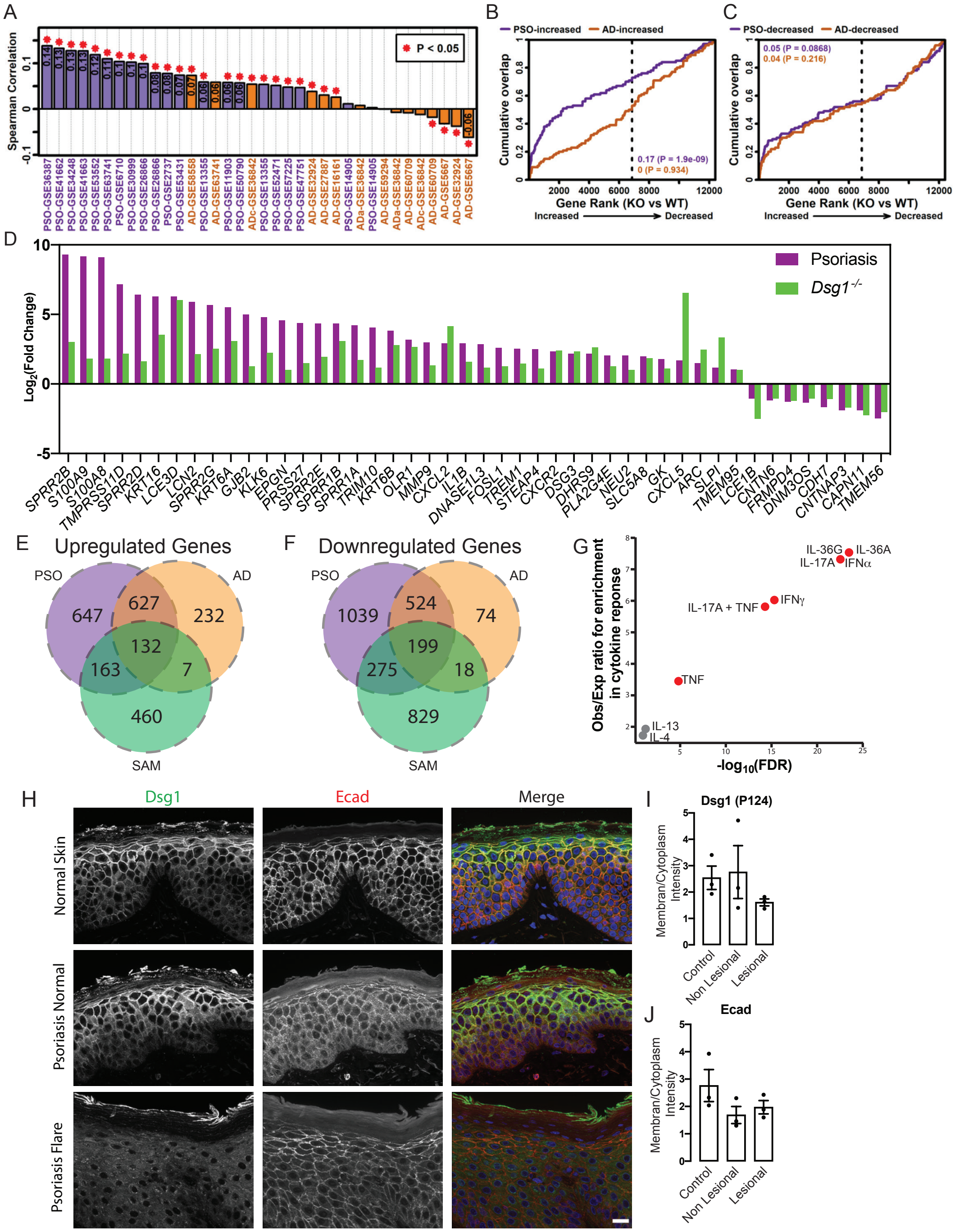


**Supplemental Figure 6. *Dsg1$^{-/-}$* mouse skin and SAM patient skin resembles psoriasis more than atopic dermatitis.** A) Gene fold change signature (*Dsg1$^{-/-}$*/*Dsg1$^{+/+}$*) were compared to those obtained from 36 comparisons yielding PSO/control (n = 21) or AD/control (n = 15) gene fold change signatures (ADa = acute atopic dermatitis; ADc = chronic atopic dermatitis). The 36 comparisons are ranked based upon the Spearman correlation coefficient estimate. B) GSEA analysis of PSO/AD-increased genes. (C) GSEA analysis of PSO/AD-decreased genes. In (B) and (C), the top 100 genes most strongly increased or decreased in each disease were analyzed. The figure shows cumulative overlap of these genes (vertical axis) with genes ranked based upon *Dsg1$^{-/-}$*/*Dsg1$^{+/+}$* FC estimates (horizontal axis). The area between each curve and the diagonal is shown with corresponding p-values (Wilcoxon rank sum test). D) Fold change for significantly upregulated or downregulated genes in both psoriasis and *Dsg1$^{-/-}$* samples. E) Overlap in upregulated genes in SAM syndrome, AD, and PSO. F) Overlap in downregulated genes in SAM syndrome, AD, and PSO. G) Cytokine response gene sets in keratinocytes overlapping with upregulated genes in psoriasis patients. H) Immunostaining for Dsg1, using the ectodomain antibody P124, and Ecad in normal skin, non-lesional skin and lesional skin from psoriasis patients. Scale bar = 20 μm. I) Quantification of Dsg1 expressed as membrane intensity over cytoplasmic intensity (n = 3). J) Quantification of Ecad expressed as membrane intensity over cytoplasmic intensity (n=3).
